# SLIRP promotes autoimmune diseases by amplifying antiviral signaling via positive feedback regulation

**DOI:** 10.1101/2024.03.28.587146

**Authors:** Doyeong Ku, Yewon Yang, Youngran Park, Daesong Jang, Namseok Lee, Yong-ki Lee, Keonyong Lee, Jaeseon Lee, Yeon Bi Han, Soojin Jang, Se Rim Choi, You-Jung Ha, Yong Seok Choi, Woo-Jin Jeong, Yun Jong Lee, Kyung Jin Lee, Seunghee Cha, Yoosik Kim

**Affiliations:** Department of Chemical and Biomolecular Engineering, Korea Advanced Institute of Science and Technology (KAIST), Daejeon, 34141, Republic of Korea; Center for RNA Research, Institute of Basic Science, Seoul, 08826, Republic of Korea; School of Biological Sciences, Seoul National University, Seoul, 08826, Republic of Korea; Department of Oral and Maxillofacial Diagnostic Science, Center for Orphaned Autoimmune Disorders, University of Florida College of Dentistry, Gainesville, Florida, 32610, United States of America; R&D Institute, ORGANOIDSCIENCES Ltd., Seongnam, 13488, Republic of Korea; Department of Pathology and Translational Medicine, Seoul National University Bundang Hospital, Seongnam, 13620, Republic of Korea; Division of Rheumatology, Department of Internal Medicine, Seoul National University Bundang Hospital, Seongnam, 13620, Republic of Korea; Medical Science Research Institute, Seoul National University Bundang Hospital, Seongnam, 13620, Republic of Korea; Department of Otorhinolaryngology - Head & Neck Surgery, Seoul National University Bundang Hospital, Seongnam, 13620, Republic of Korea; Sensory Organ Research Institute, Seoul National University Medical Research Center, Seoul National University College of Medicine, Seoul, Republic of Korea; Graduate School of Engineering Biology, KAIST, Daejeon, 34141, Republic of Korea; KAIST Institute for BioCentury (KIB), Daejeon, 34141, Republic of Korea; KAIST Institute for Health Science and Technology (KIHST), Daejeon 34141, Republic of Korea

**Keywords:** Double-stranded RNAs, antiviral signaling, mitochondrial RNAs, SLIRP, interferon response, autoimmune disease, viral infection, mitochondrial-nuclear communication, innate immune response, Sjögren’s disease

## Abstract

The abnormal innate immune response is a prominent feature underlying autoimmune diseases. One emerging factor that can trigger dysregulated immune activation is cytosolic mitochondrial double-stranded RNAs (mt-dsRNAs). However, the mechanism by which mt-dsRNAs stimulate immune responses remains poorly understood. Here, we discover SRA stem-loop interacting RNA binding protein (SLIRP) as a key amplifier of mt-dsRNA-triggered antiviral signals. In autoimmune diseases, SLIRP is commonly upregulated, and targeted knockdown of SLIRP dampens the interferon response. We find that the activation of melanoma differentiation-associated gene 5 (MDA5) by exogenous dsRNAs upregulates SLIRP, which then stabilizes mt-dsRNAs and promotes their cytosolic release to activate MDA5 further, augmenting the interferon response. Furthermore, the downregulation of SLIRP partially rescues the abnormal interferon-stimulated gene expression in autoimmune patients’ primary cells and makes cells vulnerable to certain viral infections. Our study unveils SLIRP as a pivotal mediator of interferon response through positive feedback amplification of antiviral signaling.

## INTRODUCTION

Autoimmune diseases are characterized by aberrant activation of immune response in which the body’s immune system attacks its own tissues, leading to chronic inflammation and tissue damage^1,2^. Currently, the etiology of autoimmune disease is not yet fully understood, limiting appropriate diagnosis and treatment. The dysregulated immune activation can arise from the immune system’s inability to distinguish between self and non-self signatures^3,4^. One representative non-self signature that can activate immune response is long double-stranded RNAs (dsRNAs), generally produced during viral replication^5,6^. These molecules act as pathogen-associated molecular patterns (PAMPs) and are recognized by dsRNA-specific pattern recognition receptors (PRRs), such as Toll-like receptor 3 (TLR3)^7^, retinoic acid-inducible gene-I (RIG-I), melanoma differentiation-associated gene 5 (MDA5)^8^, and protein kinase R (PKR)^9,10^. When activated by dsRNAs, these sensors trigger the expression of type I interferon (IFN) and IFN-stimulated genes (ISGs) to interfere with viral replication and suppress global translation to induce apoptosis of infected cells^11,12^.

Endogenous dsRNAs that are naturally present in our cells can also be perceived as non-self signatures, linking them to diverse autoimmune diseases^13^. Due to their immunogenic potential, the expression and recognition of dsRNAs are strictly regulated^13–15^. The dsRNAs encoded by the nuclear genome are mostly suppressed through epigenetic modification^16^. When transcribed, these RNAs are often edited by adenosine deaminase acting on RNA (ADAR), which disrupts the secondary structure to avoid recognition by dsRNA sensors^17–21^. More recently, studies showed that cells express circular RNAs with short hairpin structures to prevent activation of PKR^22,23^. However, our understanding of the regulation of mitochondrial dsRNAs (mt-dsRNAs) is highly limited.

mt-dsRNAs are generated due to bidirectional transcription of the mitochondrial circular genome^24,25^, which produces long complementary RNAs denoted as heavy and light strand RNAs^20,26–31^. Under stress conditions, mt-dsRNAs are released to the cytosol through BCL-2-associated X protein (BAX)/BCL-2- homologous antagonist/killer (BAK1) channel ^28,32,33^ where they activate MDA5 and PKR to initiate IFN response and apoptosis^28,34–36^. To limit mt-dsRNA expression, light strand mtRNAs are quickly degraded by mtRNA degradosome complex consisting of human suppressor of var1,3-like (hSUV3)^37^ and polyribonucleotide nucleotidyltransferase 1 (PNPT1)^38–40^.

Recently, studies reported misregulation of mt-dsRNAs in diseases that accompany aberrant immune activation, such as osteoarthritis (OA)^41^, Huntington’s disease^42^, alcohol-associated liver disease^43^, and autoimmune Sjögren’s disease (SjD)^44^. In particular, mt-dsRNAs are upregulated in saliva, tear, and primary salivary gland epithelial cells of SjD patients as well as in the SjD mouse model^44^. Notably, the upregulation of mtRNA expression is not limited to these inflammatory diseases. In response to viral infections, such as respiratory syncytial virus (RSV), severe acute respiratory syndrome coronavirus (SARS-CoV), and SARS-CoV-2 ^45,46^, mtRNA levels in the infected cells are elevated, suggesting a potential role of mtRNA upregulation in regulating antiviral signaling. Indeed, reducing mtRNA expression decreased the IFN response to exogenous dsRNAs and alleviated molecular phenotypes of SjD^44^. Despite the general occurrence of mtRNA upregulation and its regulatory potential on antiviral response, the underlying mechanism of mtRNA regulation remains unknown.

In this study, we investigated the regulatory mechanism and biological significance of mt-dsRNA regulation in driving aberrant immune activation. We found that mtRNA stabilizing factors, such as SRA stem-loop interacting RNA binding protein (SLIRP), are commonly upregulated in several autoimmune diseases, and targeted downregulation of SLIRP significantly reduced the IFN signature. We further analyzed the SLIRP expression, localization, and interaction with mtRNAs to elucidate the molecular mechanism behind SLIRP-mediated IFN regulation. Moreover, we examined the clinical implications by investigating the role of SLIRP in the context of SjD and viral infection. Collectively, our study establishes SLIRP as a pivotal mediator of antiviral signaling and a potential target to alleviate aberrant immune activation.

## RESULTS

### SLIRP is upregulated in autoimmune patients and regulates interferon response

Previously, we have reported that mt-dsRNA expression is upregulated in autoimmune patients^41,44^. Yet, the molecular mechanism and the regulator of mt-dsRNA expression remain unknown. Studies have established that post-transcriptional regulation of mtRNAs is a balance between degradation by RNA exonuclease 2 (REXO2)^47^ and mitochondrial degradosome complex, consisting of hSUV3^37^ and PNPT1^38–40^, and stabilization by leucine-rich pentatricopeptide repeat-containing protein (LRPPRC)^48^, SLIRP^49–51^, and mitochondrial poly(A) polymerase (MTPAP)^52^. Thus, we began our investigation by examining the expression pattern of six key regulators of mtRNA stability in autoimmune patients.

First, we reanalyzed previously published bulk RNA-seq data of monocytes of SjD and systemic lupus erythematosus (SLE) patients^53^. To determine differentially expressed genes (DEGs), gene expression in CD14+ monocytes isolated from whole blood of SjD and SLE patients (females, ages 32 to 62) were compared to those from healthy controls (HCs, females, ages 30 and 33). Interestingly, we found a significant increase in *SLIRP* mRNA expression in both SjD and SLE patients (Figure 1A), suggesting the potential association between *SLIRP* and these autoimmune diseases. At the same time, *LRPPRC*, *REXO2*, and *hSUV3* expression did not show significant differences. Of note, SjD patients exhibited a significantly elevated expression of *PNPT1*, and SLE patients showed decreased expression of *MTPAP*. However, since the upregulation of *PNPT1* and downregulation of *MTPAP* should lead to mt-dsRNA downregulation, it cannot account for the reported elevation of mtRNA expression. In addition, we reanalyzed the expression of the mtRNA regulators in other autoimmune diseases, including peripheral blood mononuclear cells (PBMCs) of rheumatoid arthritis (RA)^54^, extracellular vesicles (EVs) of secondary progressive multiple sclerosis (SPMS)^55^, and fibroblast of Aicardi-Goutières Syndrome (AGS)^56^ (Figure 1B). Notably, *SLIRP* was upregulated in all three autoimmune diseases, along with degrading factors such as *hSUV3* and *PNPT1*. Other stabilizing factors, such as *LRPPRC* and *MTPAP*, and degrading factor *REXO2* exhibited varying expression patterns across the three diseases. Given that *SLIRP* is the only upregulated regulator capable of promoting mtRNA expression, we focused on SLIRP.

**Figure 1.**
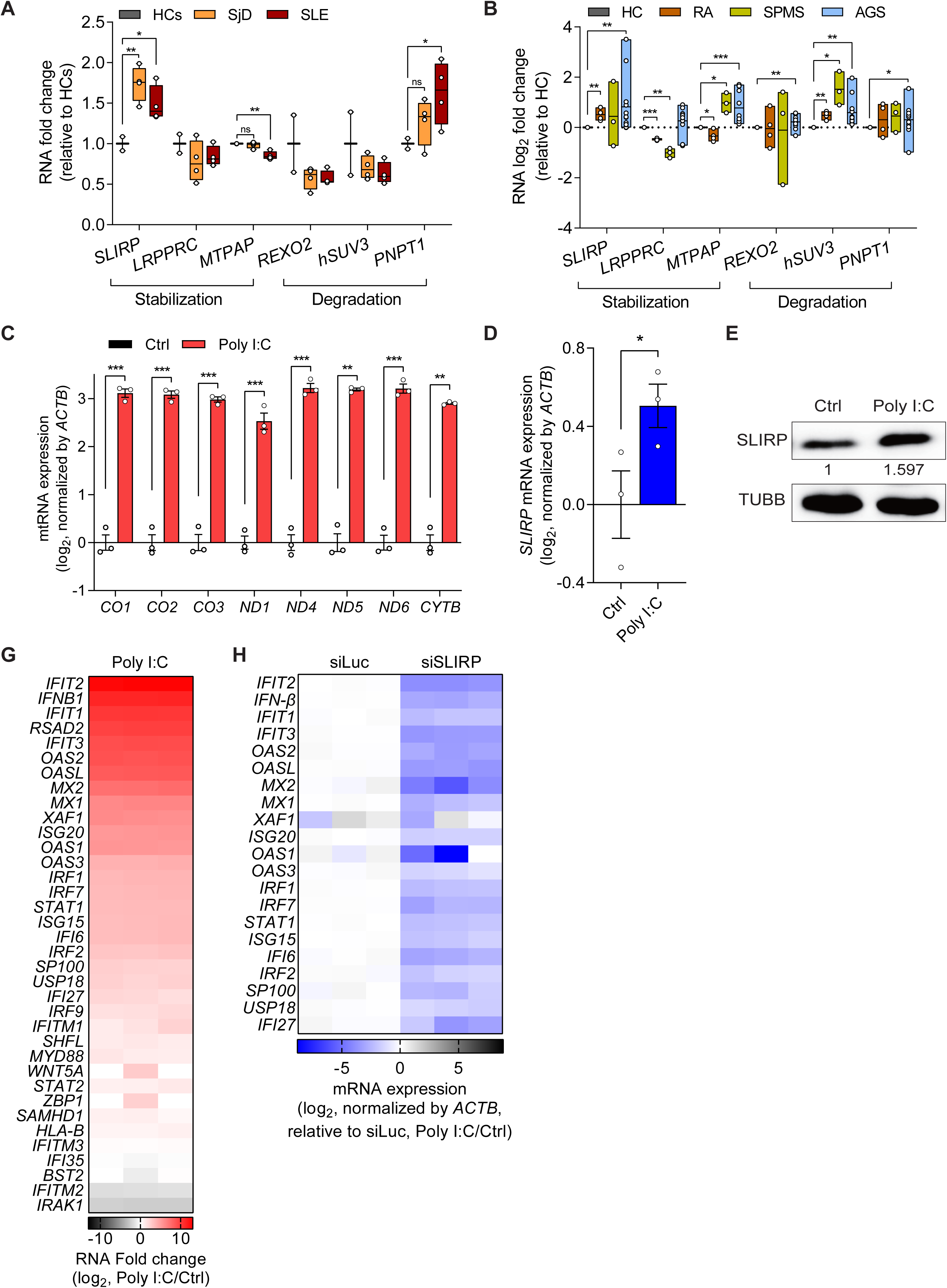
SLIRP is upregulated and its downregulation regulates ISG induction in autoimmune diseases. (A) Analysis of gene expression related to mtRNA stability in monocytes of HCs (n=2), SjD (n=4), and SLE (n=4) patients. (B) Log_2_ fold change for genes related to mtRNA stability in RA (n=4), SPMS (n=3), and AGS (n=12) patients. For (A) and (B), reanalyses of publicly available RNA-seq data are shown. (C) Analysis of total mtRNA expression upon poly I:C transfection. (D, E) Analysis of *SLIRP* mRNA (D) and protein (E) expression in HCT116 cells transfected with poly I:C. (F) The GO analysis of DEGs of poly I:C-transfected cells. Genes with log_2_ fold change over 2 and p-value less than 0.05 were analyzed. (G) Heatmap of log_2_ fold changes for type I IFN genes upon poly I:C transfection. The three columns represent biological replicates. (H) Heatmap of log2 fold change of type I IFN genes upon poly I:C transfection in control and SLIRP-deficient cells. The three columns represent biological replicates. Unless mentioned, three independent experiments were carried out, and error bars denote s.e.m. All of the statistical significances were calculated using one-tail Student’s t-tests; * p ≤ 0.05, ** p ≤ 0.01, and *** p ≤ 0.001.

To further investigate the significance of *SLIRP* upregulation, we employed polyinosinic-polycyticylic acid (poly I:C), a synthesized double-stranded RNA widely utilized to induce the inflammatory responses that can mimic etiology of autoimmune diseases^44,57–61^. First, we confirmed that stimulating cells with poly I:C results in significant upregulation of mtRNAs (Figure 1C). Moreover, consistent with patient data, we found about a 50% increase in both mRNA and protein expression of SLIRP in poly I:C-transfected cells (Figures 1D and 1E). To test whether SLIRP can mediate the downstream IFN response to poly I:C, we performed transcriptome analysis and identified a list of IFNs and ISGs that are strongly induced by poly I:C (Figure 1G). Interestingly, we found that SLIRP downregulation resulted in a significant reduction of all of them (Figure 1H). Of note, these genes are still induced by poly I:C in SLIRP-deficient cells, but the degree of induction was significantly attenuated compared to the control cells. Collectively, we suggest SLIRP as a potential regulator in modulating aberrant IFN response in autoimmune diseases.

### SLIRP regulates IFN response by stabilizing mtRNAs

SLIRP is an mtRNA binding protein that interacts with stem-loop structures in the 3′ untranslated regions to protect the RNA from decay^49–51^. Therefore, we asked whether SLIRP can affect the IFN response by stabilizing and elevating mtRNA expression during antiviral signaling. First, we investigated whether mtRNA induction is a general consequence of antiviral signaling to dsRNAs or specific to poly I:C. We found that transfection of long synthetic dsRNAs and poly I:C resulted in the strong upregulation of both total mtRNAs and mt-dsRNAs (Figures 2A and 2B). Of note, mt-dsRNA expression was inferred by examining the individual expression of heavy and their complementary light strand mtRNAs via strand-specific reverse transcription^62^. Transfection of short synthetic dsRNAs (79 bp duplex RNA with random sequences) increased the expression of light strand mtRNAs, with no notable alteration in the heavy strands and total mtRNAs. Meanwhile, transfection of long synthetic dsRNAs (717 bp duplex RNA derived from *EGFP* sequences) and poly I:C (average length of 1,000∼1,800 bp) increased mtRNA expression from both strands. Interestingly, long synthetic dsRNAs and poly I:C yielded similar effects on mtRNAs, suggesting that the upregulation of mtRNAs is not dsRNA-sequence specific but rather a general downstream response to long dsRNA transfection.

**Figure 2.**
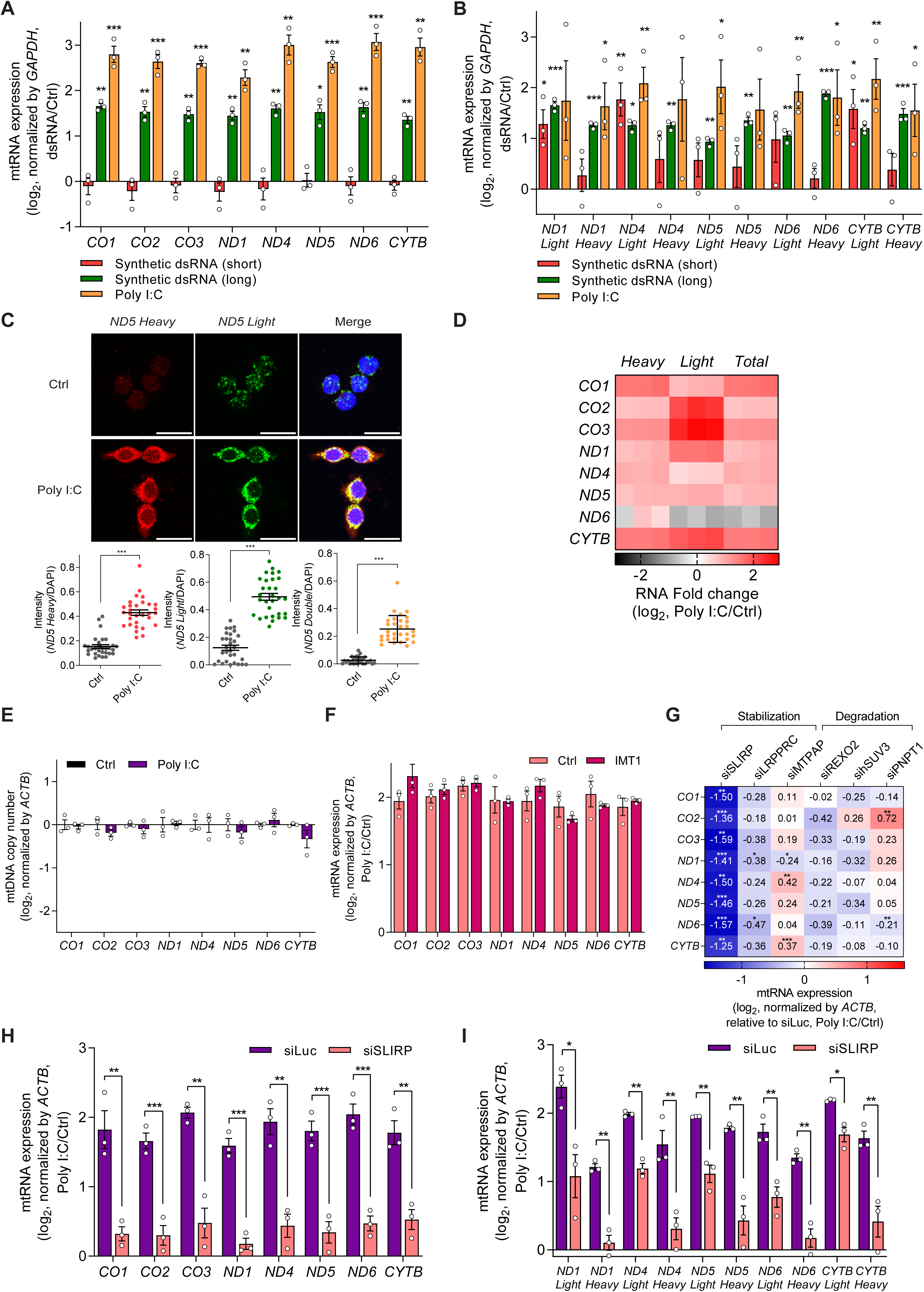
Stabilization by SLIRP results in mtRNA upregulation during antiviral response. (A, B) Analysis of total (A) and strand-specific (B) mtRNA expression upon introduction of dsRNAs with different lengths in HCT116 cells. (C) smRNA-FISH analysis for the individual strand of *ND5* mtRNAs in poly I:C transfected cells. Nuclei were stained using DAPI. Quantified signal intensities calculated by taking an average of over 30 images are presented below. Scale bar, 25 μm. (D) Heatmap of log_2_ fold changes for the two strands of mtRNAs upon poly I:C transfection. The three columns represent biological replicates. (E) Analysis of mtDNA copy number upon poly I:C transfection in HCT116 cells. (F) Normalized total mtRNA levels upon poly I:C transfection 48 h after IMT1 treatment. Data are normalized only to those of DEPC-transfected cells in order to analyze the degree of mtRNA induction by poly I:C. (G) Heatmap of the degree of mtRNA induction upon poly I:C transfection in cells transfected with the indicated siRNAs. (H, I) Analysis of total (H) and strand-specific (I) mtRNAs in control and SLIRP knocked-down HCT116 cells. Three independent experiments were carried out, and error bars denote s.e.m. All of the statistical significances were calculated using one-tail Student’s t-tests; * p ≤ 0.05, ** p ≤ 0.01, and *** p ≤ 0.001.

Through single-molecule RNA fluorescent in situ hybridization (smRNA-FISH), we visualized and quantified the increased expression of *ND5* mtRNAs upon poly I:C transfection (Figure 2C). Of note, *ND5* was chosen as a representative mtRNA because *ND5* light and heavy strand RNAs showed strong induction in our current and previous studies^41,44,62^. To complement smRNA-FISH data, we performed RNA-seq analysis and found that both strands of all examined mtRNAs, except for the light strand *ND6*, were upregulated upon poly I:C transfection (Figure 2D). Such upregulation of mtRNAs was consistently observed across multiple cell lines, such as A549 lung adenocarcinoma and PC3 prostatic adenocarcinoma, indicating that it might be a common downstream response to dsRNA-mediated antiviral response (Figures S1A and S1B).

To investigate the molecular mechanism behind the mtRNA upregulation, we first analyzed and confirmed that the mtDNA replication, as determined by qPCR and gel electrophoresis, is not affected by poly I:C (Figure 2E and S2A). To assess the effect of mtRNA transcription, we pre-treated cells with an inhibitor of mitochondrial transcription (inositol 4-methyltransferase, IMT1)^63^ for 41 h prior to poly I:C transfection, which resulted in a significant decrease in mtRNA levels (Figure S2B). Yet, despite the inhibition of mtRNA transcription, we still observed clear increase in mtRNA expression upon poly I:C transfection (Figure 2F). The similar degree of induction between mock and IMT1-treated cells ruled out the possibility of leaky transcription due to imperfect inhibition of mtRNA polymerase by IMT1. Of note, we normalized mtRNA expression prior to poly I:C transfection to focus on the degree of induction rather than the relative expression level of mtRNAs upon poly I:C transfection.

To test whether mtRNA stability was responsible for the increased mtRNA expression, we downregulated the six key regulators involved in mtRNA stability individually using siRNAs and examined mtRNA induction by poly I:C. To focus on the degree of induction, we normalized the mtRNA expression in cells deficient in the individual target gene without poly I:C transfection. Among these candidates, we found that SLIRP knockdown resulted in a significant decrease in the degree of induction for all mtRNAs examined (Figures 2G and S2C-G). More importantly, in SLIRP-deficient cells, the mtRNA induction was nearly completely abolished, indicating that SLIRP is the primary factor in upregulating mtRNA expression during antiviral response to exogenous dsRNAs (Figure 2H). When we performed strand-specific RT-qPCR to examine the two strands of mt-dsRNAs independently, we found that the SLIRP knockdown decreased induction in both strands, with a stronger effect on the heavy strand mtRNAs (Figure 2I).

Next, we asked about the generality of the observed SLIRP-mediated upregulation of mtRNAs during innate immune response in general. To test, we obtained several representative innate immune stressors based on their well-known non-self signatures and interacting PRRs and evaluated their impact on mtRNA expression (Figure S3A). We used lipopolysaccharide (LPS) stimulating TLR4 as a non-nucleic acid PAMP, single-strand poly-uridine (ssPolyU) as an agonist of TLR7 and TLR8, 5′ triphosphate hairpin RNA (3p-hpRNA) as an agonist of RIG-I, and synthetic dsRNAs (short, long, and poly I:C) as agonists of RIG-I, MDA5, TLR3, and PKR. We also examined the DNA stressor, CpG oligodeoxynucleotides (CpG ODN), which can act as an agonist of cyclic GMP-AMP synthase (cGAS) and TLR9. In contrast to dsRNAs, LPS treatment did not cause a significant change in total mtRNA expression, while ssPolyU treatment and 3p-hpRNA transfection even reduced total mtRNA expression (Figures S3B-D). Treatment with poly I:C to activate TLR3 resulted in a slight increase in total mtRNA expression with a large variance between biological replicates (Figure S3E). Lastly, neither treatment nor transfection of CpG ODN to activate TLR9 or cGAS, respectively, elicited any significant alterations in total mtRNA expression (Figures S3F and S3G). Therefore, mtRNA induction occurred most strongly as a downstream response to long dsRNA transfection, and we focused on poly I:C for the remainder of the study.

### Mitochondrial localization of SLIRP is enhanced by poly I:C

Our data suggests a model where the RNA stabilizing effect of SLIRP is enhanced during antiviral response to exogenous dsRNAs and results in increased mtRNA levels. When we examined the SLIRP subcellular localization, we found increased SLIRP expression in the membrane fraction, which contains mitochondria (Figure 3A). Indeed, SLIRP showed a higher degree of colocalization with the mitochondria upon poly I:C transfection (Figure 3B). For more direct confirmation, we employed formaldehyde crosslinking and immunoprecipitation (fCLIP) and examined SLIRP-mtRNA interactions (Figure S4A). In cells transfected with poly I:C, SLIRP was more strongly bound to mtRNAs, except for *ND5*, which showed large variability (Figure 3C). Moreover, we analyzed the mtRNA stability in control and SLIRP-deficient cells after blocking mtRNA transcription using IMT1. Consistent with our earlier data, we found a dramatic increase in mtRNA stability only in poly I:C-transfected cells (Figures 3D and S4B). Interestingly, we no longer observed increased mtRNA stability by poly I:C in SLIRP-deficient cells (Figures 3D and S4B). There was a slight increase in mtRNA levels in 2 h sample, but the magnitude was minor compared to that of the control (siLuc) cells.

**Figure 3.**
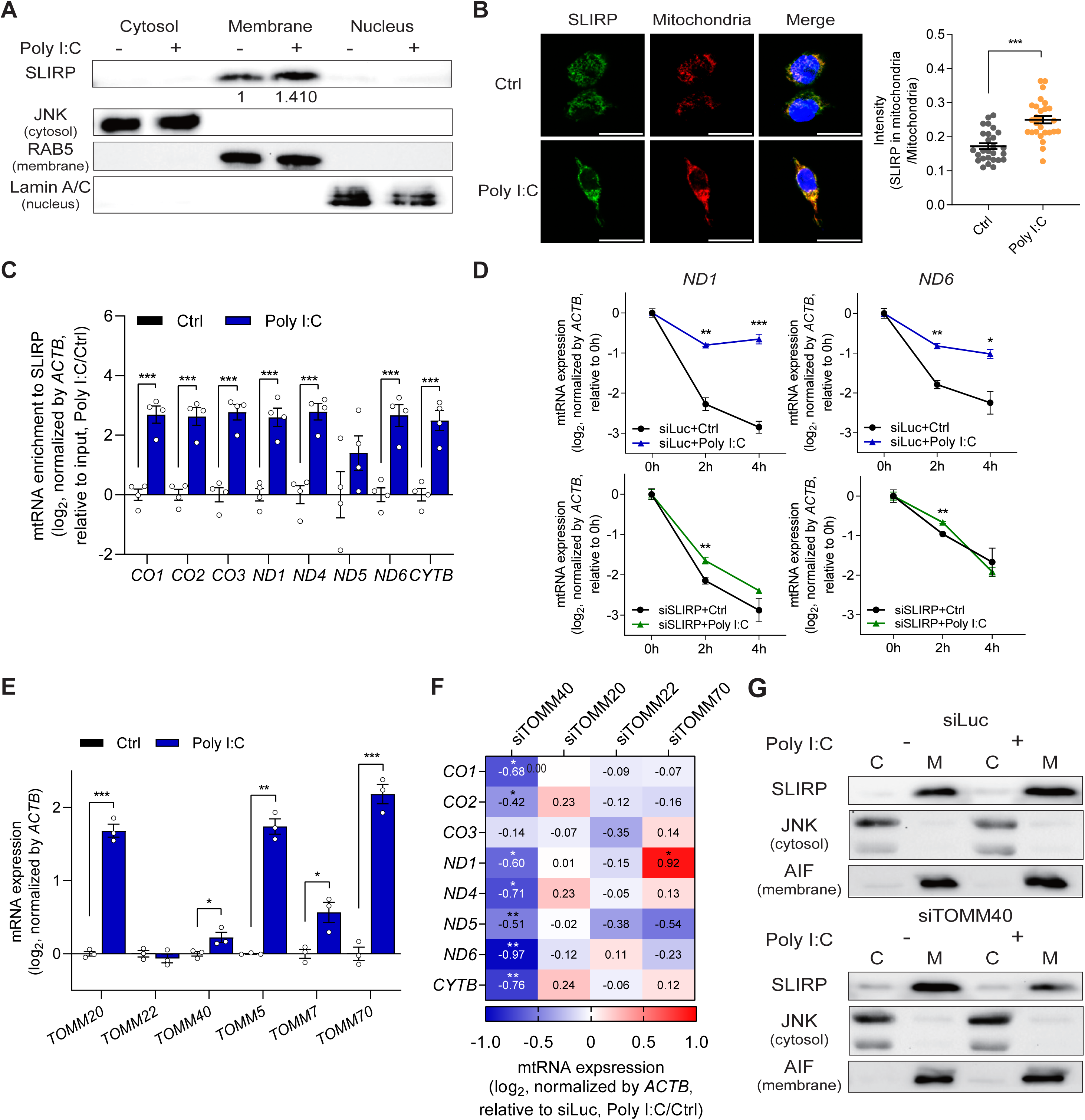
Mitochondrial import of SLIRP is upregulated during antiviral response. (A) The expression of SLIRP in free cytosol, membrane organelle, and nucleus upon poly I:C transfection. The following proteins are used to confirm successful subcellular fractionation: JNK for the free cytosol, RAB5 for the membrane, and Lamin A/C for the nucleus. (B) Representative fluorescence images of SLIRP and mitochondria upon poly I:C transfection. Scale bar, 25 μm. Quantified signal intensities averaged over 17 images are presented below. (C) Interaction between SLIRP and mtRNAs upon poly I:C transfection was examined by SLIRP fCLIP followed by RT-qPCR analysis (n=4). (D) Analysis of mtRNA stability before and after poly I:C transfection in control and SLIRP-deficient cells. (E) Analysis of the RNA expression of genes involved in the TOM complex upon poly I:C stimulation. (F) Heatmap showing the degree of mtRNA induction by poly I:C after downregulating the indicated target genes involved in the TOM complex. (G) Western blot analysis for SLIRP expression upon poly I:C in the cytosol and membrane fraction, with or without TOMM40. JNK is used as a marker for the cytosol fraction while AIF is used as a marker for the membrane fraction. Unless mentioned, three independent experiments were carried out, and error bars denote s.e.m. All of the statistical significances were calculated using one-tail Student’s t-tests; * p ≤ 0.05, ** p ≤ 0.01, and *** p ≤ 0.001.

We further investigated the mechanism responsible for increased SLIRP localization to mitochondria during antiviral response to poly I:C. To this end, we examined the translocase of the outer membrane (TOM) complex, which plays a crucial role in recognizing mitochondrial targeting signals to facilitate the entry of precursor proteins into the mitochondria^64–66^. The key components of the TOM complex include translocase of outer mitochondrial membrane 20 (TOMM20) and TOMM22^67,68^, the central channel protein, TOMM40^69^, and an accessory protein TOMM70^70^. We examined the effect of poly I:C transfection on their mRNA expression and found increased expression in all except for *TOMM22* (Figure 3E). We then applied a similar approach as above and knocked-down these four *TOMM* genes individually using siRNAs and examined mtRNA induction by poly I:C. Remarkably, our mini-screening revealed that knockdown of TOMM40, resulted in decreased induction of mtRNAs upon poly I:C stimulation, except for *CO3* (Figure 3F and S5A-D). To test whether the knockdown of TOMM40 affected SLIRP localization, we performed subcellular fractionation and examined SLIRP expression. While control cells showed increased SLIRP localization to the membrane fraction upon poly I:C transfection, in TOMM40-deficient cells, SLIRP recruitment to the membrane fraction was not facilitated (Figure 3G). Our data indicate that the increased expression and enhanced mitochondrial import of SLIRP by the mitochondrial protein import system, particularly TOMM40, plays a pivotal role in the induction of mtRNA via increased stability during antiviral signaling.

### SLIRP-mtRNA-MDA5 positive feedback loop amplifies antiviral signaling

When present in the cytosol, mt-dsRNAs act as self-immunogens and activate PRRs, activating antiviral signaling. To investigate how SLIRP-mediated mtRNA stabilization contributes to ISG expression, we examined the role of SLIRP in the cytosolic release of mtRNAs. We found that SLIRP-deficient cells showed reduced cytosolic levels for most mtRNAs in response to poly I:C (Figure S6A and 4A). Since mtRNAs and mt-dsRNAs are released to the cytosol through BAX/BAK micropores^28,32,33^, we downregulated BAK1 to hinder the cytosolic release of mtRNAs and examined the expression of selected ISGs. All examined ISGs showed decreased expression in BAK1 knocked-down cells (Figure 4B). To our surprise, we found that the induction of most mtRNAs upon poly I:C stimulation was also significantly attenuated in BAK1-deficient cells (Figure 4C), indicating that the cytosolic release of mtRNAs might contribute to upregulation of their own expression. This creates a positive feedback loop where activation of antiviral signaling by exogenous dsRNAs upregulates SLIRP and facilitates the mitochondrial import of SLIRP, where it stabilizes mtRNAs, which are then released to the cytosol to amplify the antiviral signaling further.

**Figure 4.**
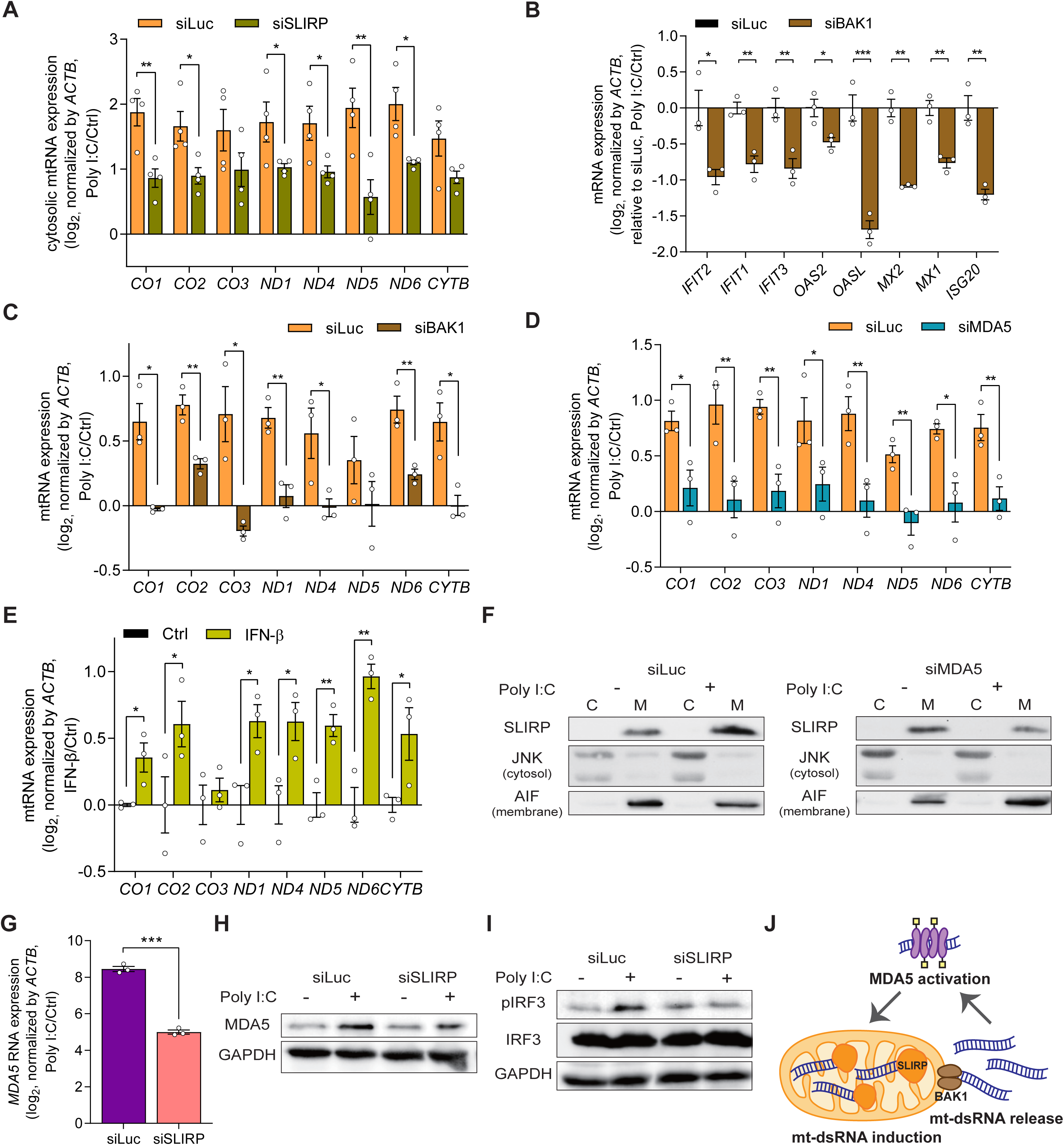
Antiviral signaling is amplified via the MDA5-mtRNA-SLIRP positive feedback loop. (A) Cytosolic mtRNA expression in poly I:C transfected cells with or without SLIRP (n=4). (B, C) The effect of BAK1 downregulation on the ISG (B) and mtRNA (C) induction by poly I:C. (D) Analysis of mtRNA expression upon poly I:C transfection with or without MDA5. (E) The mtRNA expression upon IFN-β treatment. (F) Western blot analysis for SLIRP localization upon poly I:C with or without MDA5. JNK is used as a marker for the cytosol fraction while AIF is used as a marker for the membrane fraction. (G, H) The expression of *MDA5* mRNA (G) and protein (H) expression after poly I:C transfection in cells deficient in SLIRP. (I) The effect of SLIRP downregulation on the IRF3 phosphorylation upon poly I:C. (J) A model for the positive feedback amplification of antiviral signaling consists of mtRNA induction, cytosolic release, and MDA5 activation. Unless mentioned, three independent experiments were carried out, and error bars denote s.e.m. All of the statistical significances were calculated using one-tail Student’s t-tests; * p ≤ 0.05, ** p ≤ 0.01, and *** p ≤ 0.001.

A missing component in our proposed feedback regulation is the factor that mediates SLIRP and mtRNA induction upon poly I:C transfection. To test, we downregulated the dsRNA sensors individually and examined mtRNA levels. Interestingly, MDA5 knockdown resulted in a significantly reduced induction of mtRNAs upon poly I:C transfection (Figures 4D and S6B-D). In the presence of long dsRNAs, MDA5 forms a filament along the dsRNA, recruits mitochondrial antiviral signaling protein, and phosphorylates interferon regulatory factor 3 (IRF3) to trigger the type I IFN response, including IFN-β^71^. Indeed, treating cells with IFN-β was sufficient in upregulating mtRNA expression even without poly I:C stimulation (Figure 4E). Moreover, in MDA5-deficient cells, SLIRP localization to mitochondria was no longer enhanced (Figure 4F). This suggests that MDA5 and the downstream IFN-β signaling are responsible for the mtRNA induction.

MDA5 is one of the ISGs and upregulates its own expression during antiviral response. Considering that SLIRP affects the global induction of ISGs during dsRNA stress, we examined whether MDA5 expression is under the control of SLIRP as well. Notably, we found that SLIRP knockdown results in a dramatic decrease in *MDA5* mRNA and protein expression (Figures 4G and 4H). The decreased MDA5 expression eventually resulted in the attenuation of antiviral signaling, as the phosphor-IRF3 (pIRF3) was significantly reduced in SLIRP knocked-down cells (Figure 4I). Collectively, we establish a positive feedback model of antiviral signaling consisting of MDA5 activation, enhanced SLIRP mitochondrial localization, stabilization and cytosolic release of mtRNAs and mt-dsRNAs, and further activation of MDA5 (Figure 4J).

### SLIRP can drive abnormal IFN signatures in SjD patients

Next, we examined the significance of SLIRP as a putative regulator of aberrant antiviral signaling in autoimmune disease. Among various autoimmune diseases that showed elevated SLIRP expression, we focused on SjD as mt-dsRNAs are involved in the pathogenesis of SjD^44^. First, we investigated the upregulation of SLIRP in different types of SjD. We obtained and analyzed monocytes from SjD patients of varying ages, including five adults (SjD) and four children diagnosed with SjD (cSjD) as well as monocytes from various control groups: five HCs (mostly young adults), four symptomatic pediatric patients who do not meet the criteria for cSjD diagnosis (Non-cSjD), and six non-cSjD patients with the potential to develop SjD in the future based on their positive lip biopsy results (BxP). Our transcriptome analysis revealed an overall increase in *SLIRP* mRNA expression in SjD patients, except for one patient, compared to the control groups, indicating a potential association between *SLIRP* and SjD. Two cSjD patients also showed increased SLIRP expression, while the others did not. In contrast, other mtRNA stabilizing factors, such as *LRPPRC* and *MTPAP*, did not show significant changes in their expression among SjD patients and control groups (Figure 5A). When we examined mtRNA degradosome components, *PNPT1* was significantly increased in SjD patients, which cannot account for the observed increase in mtRNAs (Figure 5B). The expression of other genes related to mtRNA degradosome in SjD and cSjD was indistinguishable from those of the control groups (Figure 5B).

**Figure 5.**
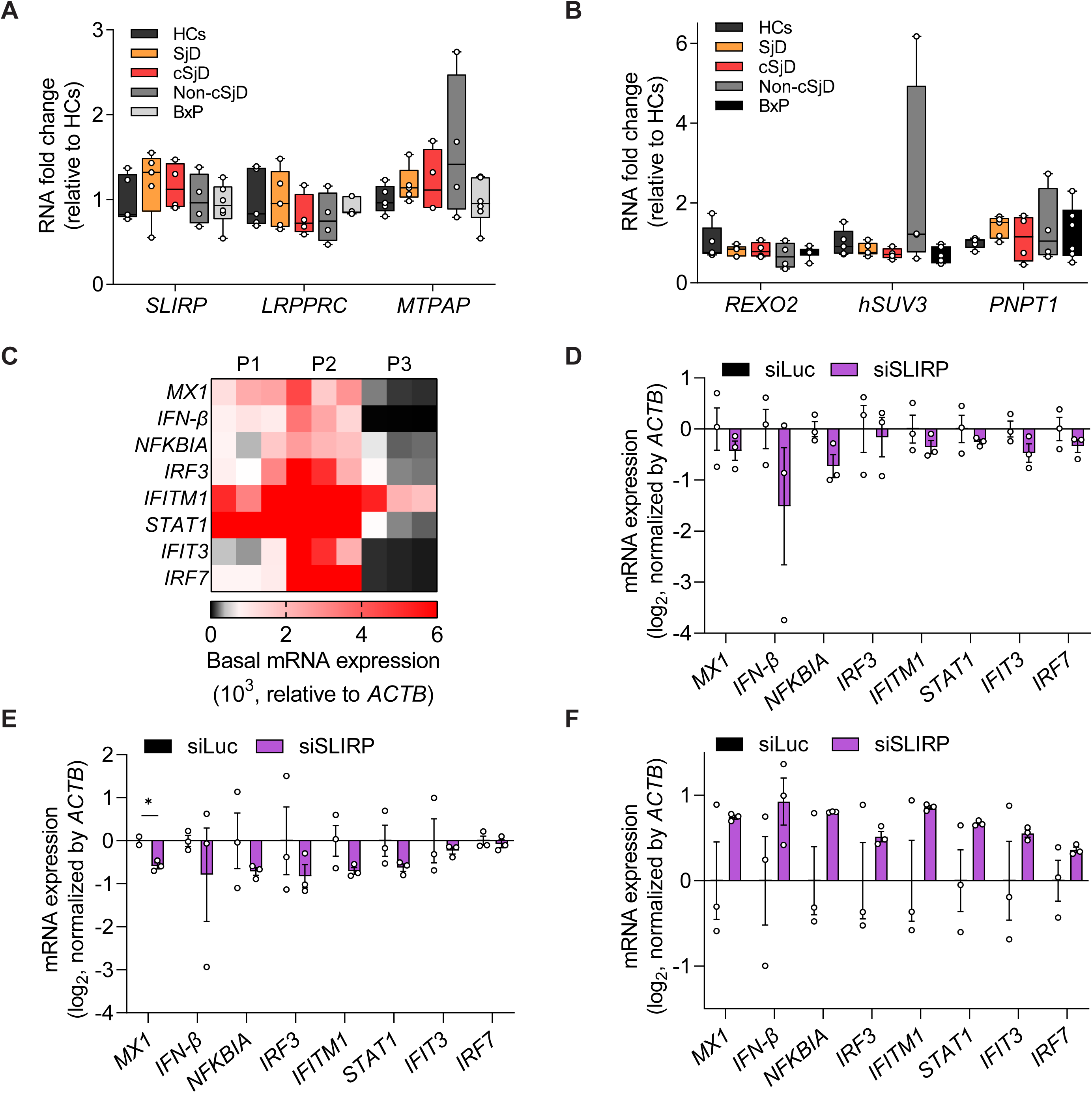
SLIRP is associated with the immune signature of SjD and SLE patients. (A, B) Analysis of gene expression related to mtRNA stability in monocytes of HCs (n=5), SjD (n=5), cSjD (n=4), Non-cSjD (n=4), and BxP (n=6). (C) The heatmap representing the unstimulated expression of selected ISGs in primary minor salivary gland cells isolated from three SjD patients. The three columns represent three biological replicates. (D-F) Analysis of ISG expression after knockdown of SLIRP in primary minor salivary gland cells of three patients from (C). Unless mentioned, n=3 and error bars denote s.e.m for all except for the organoid data. All of the statistical significances were calculated using one-tail Student’s t-tests; * p ≤ 0.05, ** p ≤ 0.01, and *** p ≤ 0.001.

We further investigate the role of SLIRP in aberrant IFN signatures in SjD by analyzing the primary cells derived from patients’ minor salivary gland lip biopsy specimens. We first characterized the IFN signature and found that two patients (P1 and P2) exhibited elevated expression of *IFN-*β and several ISGs related to SjD, whereas their downregulation was noted in P3 (Figure 6C). Interestingly, the further downregulation of SLIRP in patients’ primary cells resulted in reduced *IFN-*β and some ISG expression for both P1 and P2 (Figures 6D and 6E), albeit with large variations. On the other hand, SLIRP downregulation in cells from P3 increased *IFN-* β and ISG expression (Figure 6F).

**Figure 6.**
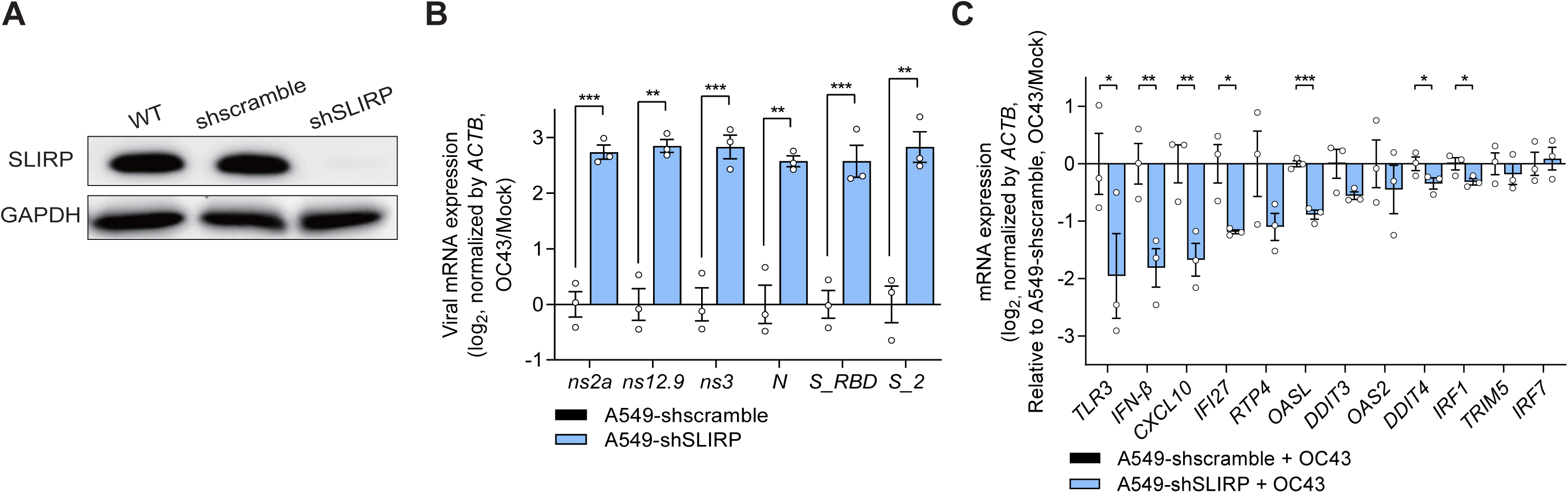
SLIRP downregulation enhances OC43 infection and attenuates ISG induction. (A) Western blot analysis of SLIRP expression in A549 cells transduced with shSLIRP. (B, C) Analysis of viral mRNA expression (B) and ISG induction (C) after infecting SLIRP-deficient cells with OC43 at MOI = 1. N=3 and error bars denote s.e.m. All of the statistical significances were calculated using one-tail Student’s t-tests; * p ≤ 0.05, ** p ≤ 0.01, and *** p ≤ 0.001.

### SLIRP mediates cellular defense against human coronavirus OC43

Considering that poly I:C is commonly used to mimic viral infection, we investigated the relevance of mtRNA stabilization by SLIRP in viral infection. To test, we knocked-down SLIRP in A549 cells using shRNAs and infected cells with human coronavirus OC43 or human influenza A virus (A/Puerto Rico/8/1934(H1N1), PR8) (Figure 6A). OC43 is a positive-strand RNA virus known to produce dsRNAs in cells during viral replication^72^, whereas PR8 is a negative-strand RNA virus with low dsRNA generation^5^. We found that the viral mRNA expression of OC43 was significantly increased in SLIRP-deficient cells, indicating enhanced viral replication (Figure 6B). Interestingly, despite the increased viral replication, the expression of *IFN-*β and selected ISGs was lower, indicating impaired innate immune response in SLIRP-deficient cells (Figure 6C). When we performed a similar experiment for the PR8 virus, SLIRP downregulation decreased the level of viral mRNA (Figure S7A). In addition, the ISG expression was unaffected in SLIRP-deficient cells infected with the PR8 virus (Figure S7B).

To extend our findings to the SARS-CoV-2 virus, a positive-strand RNA virus, we infected Calu3 and A549-ACE2 cells with the SARS-CoV-2 and examined the effect of SLIRP downregulation on viral replication. We first confirmed the knockdown efficiency of SLIRP in Calu3 via western blotting (Figure S7C). We then infected Calu3 cells with SARS-CoV-2 at MOI of 1 and 0.1, but found that SLIRP downregulation did not affect viral mRNA expression (Figure S7D). The expression of *IFN-*β and *TLR3* was even increased upon SARS-CoV-2 infection in cells deficient in SLIRP (Figure S7E). Next, we used A549-ACE2 as a different cell line susceptible to SARS-CoV-2 infection. Again, we began by confirming the degree of SLIRP knockdown by shSLIRP in A549-ACE2 cells (Figure S7F) and infected cells with SARS-CoV-2 virus at MOI of 1 and 0.1. However, for A549-ACE2, SLIRP downregulation decreased viral mRNA expression (Figure S7G). Moreover, SLIRP downregulation did not affect the expression of *TLR3* and *IFI27* while the expression of *IFN-*β and *CXCL10* was increased (Figure S7H).

## DISCUSSION

Our study establishes an amplification of antiviral signaling by SLIRP through the stabilization of mt-dsRNAs, leading to a robust IFN response to exogenous dsRNAs. According to our model, activation of MDA5 by dsRNAs and downstream IFN response upregulates SLIRP and facilitates SLIRP localization to mitochondria through an enhanced mitochondrial import system, especially TOMM40. Inside mitochondria, SLIRP stabilizes mtRNAs to elevate their expression. The increased mtRNAs are then released to the cytosol via BAX/BAK micropores, where they are recognized by MDA5 and other PRRs to activate them, completing the positive feedback amplification. Moreover, our study suggests SLIRP as a putative regulator of aberrant antiviral signatures found in autoimmune patients such as SjD and a key defense molecule against viruses, including OC43.

Our study attempted to address the biological significance behind the SLIRP- mediated amplification of the antiviral response in the context of autoimmune disease and viral infection. The upregulation of SLIRP in the monocytes of SjD and SLE, together with the upregulation of mtRNAs, suggest a potential role of SLIRP in the development of autoimmune disease. Indeed, the elevation of mtRNAs is associated with the pathogenesis of SjD, while SLE patients produce auto-antibodies that recognize mtRNAs^73^. In addition, the abnormal activation of PRRs and antiviral signaling are well-known signatures of autoimmune diseases^74–78^. Moreover, our study suggests SLIRP as a potential target to alleviate the upregulation of IFN signature in autoimmune patients, as downregulation of SLIRP in the primary minor salivary gland cells of SjD patients resulted in a moderate reduction in the expression of selected ISGs. Yet, one of the patient cells with the lowest ISG expression showed no further decrease in ISG mRNA levels upon SLIRP knockdown. One possibility is that targeting SLIRP might provide benefits to patients with high ISG signatures. Although the number of patients is too few (due to the difficult nature of obtaining samples of rare SjD disease patient samples) to make a conclusive remark, our findings suggest a potential for downregulating SLIRP to attenuate abnormal IFN signatures in SjD patients who exhibited elevated IFN and ISG expression. Future investigation using additional patients with various ISG expression patterns may provide a specific patient group where SLIRP downregulation may alleviate aberrant ISG expression.

Numerous previous studies showed the association of mt-dsRNAs with aberrant immune activation during the development of inflammatory and autoimmune diseases^41–44^. Moreover, mt-dsRNAs can be released to the cytosol during oxidative stress and mitochondrial dysfunction as well as in response to DNA damage and activate PKR to initiate apoptotic programs^34,41^. In the current study, we focused on mt-dsRNA induction during antiviral response and identified key components, such as SLIRP, TOMM40, and MDA5, that are required for mtRNA induction and subsequent amplification of the IFN response. Consistent with our results, a recent proteomics study identified TOMM40 as one of six consistently upregulated proteins in SLE patients^79^. In the future, it would be interesting to study whether SLIRP is also involved in other stress conditions where mt-dsRNAs activate the cytosolic dsRNAs sensors.

Our study primarily used poly I:C to study the downstream events of antiviral signaling. Although poly I:C is commonly used to mimic viral infection, viruses develop multiple mechanisms to avoid dsRNA sensing inside the cell^80–83^. For example, certain viruses form a specific replication compartment within the cytosol to shield the viral dsRNAs from PRRs^81^. The *Flaviviridae* family of influenza viruses can replicate in the nucleus to avoid cytosolic PRRs^84^. Moreover, morbillivirus in the *Paramyxoviridae* family post-translationally modify MDA5 to prevent its activation^85^. Similarly, SARS-CoV-2 encodes ORF3c protein that blocks MDA5 activation^86^. Perhaps, these strategies allow viruses to avoid SLIRP-mediated amplification of antiviral response. Of note, in our study, the knockdown of SLIRP resulted in decreased viral replication for PR8 and SARS-CoV-2, but we believe that this reflects decreased proliferation in SLIRP knocked-down cells instead of its effect on antiviral signaling. At the same time, in the case of OC43 human coronavirus, knockdown of SLIRP resulted in decreased antiviral signaling while viral replication was enhanced. These results suggest that MDA5-SLIRP-mtRNA feedback regulation of antiviral signaling might have been developed as a mechanism for cellular defense against viruses. In this context, mitochondria, products of endosymbiosis, are working together with the host genomic factor (SLIRP) to provide immunogenic materials (mt-dsRNAs) to defend the host cell against foreign threats (viruses). It remains to be investigated how viruses evolved to avoid PRR recognition and prevent the initiation of the positive feedback loop.

In summary, our study underscores nuclear-mitochondrial communication during antiviral signaling via SLIRP, which utilizes mitochondrial immunogenic materials as a snowball for robust immune response. Detailed analysis of the mechanism and application of SLIRP-mediated mtRNA regulation will enhance our understanding of aberrant immune signatures in autoimmune diseases and host defense against viruses.

### Limitations of the study

One limitation of our study is the number of patients’ primary cells used to analyze the clinical significance of SLIRP upregulation in autoimmune diseases. Due to the rare nature of SjD occurrence, we could only analyze three patient samples, two of which showed elevated IFN signatures. Nevertheless, our data show a potential to alleviate ISG expression by targeting SLIRP for those with elevated IFN response. Another limitation of our study is the virus-specific effect of SLIRP downregulation. Knockdown of SLIRP resulted in decreased IFN response and enhanced viral replication for only OC43 human coronavirus. As discussed above, one possibility is that the MDA5-SLIRP-mtRNA regulatory axis is not invoked in other viruses like PR8 and SARS-CoV-2 as viruses developed means to evade host cells’ defense systems. Targeting these evasion mechanisms by viruses will solidify our findings that SLIRP and mtRNAs play key regulatory roles during innate immune response.

## Supporting information

Supplemental informations

## ACKNOWLEDGEMENTS

The authors thank all members of the Cha and Kim laboratories for their helpful discussions and comments on the paper. We thank Dr. V. Narry Kim from Seoul National University for providing Calu3 cells and insightful suggestions on the study. This work was supported by the Ministry of Health & Welfare grant number HI21C1501 and NIH/NIDCR DE32707.

## AUTHOR CONTRIBUTIONS

D.K., S.C., and Y.K. conceived and designed the study. D.K. conducted most of the experiments. Y.Y performed experiments from figure S4. Y.P. performed SARS-CoV-2 experiments. D.J. prepared patient’s monocytes and primary cells. K.L. and N.L. performed the bioinformatics analyses. Y.L. prepared OC43 virus experiments. J.L., Y.H., S.J., S.C., Y.H., Y.C., W.J., Y.L., and K.L. prepared patient’s samples. D.K., Y.Y., and Y.K. analyzed data, interpreted the results, and generated the figures. D.K., Y.P., D.J., Y.K. wrote the original draft. D.K., Y.Y., Y.L., V.K., K.L., S.C., and Y.K. reviewed and edited the manuscript. S.C. and Y.K. supervised the study.

## DECLARATION OF INTERESTS

Jaeseon Lee, Soojin Jang, and Kyung Jin Lee are employees of ORGANOIDSCIENCES Ltd. Other authors have no conflict of interest to declare.

## SUPPLEMENTAL INFORMATION TITLES AND LEGENDS

Document S1. Figures S1-S7 and Tables S1-S5.

## STAR Methods

### Key resources table

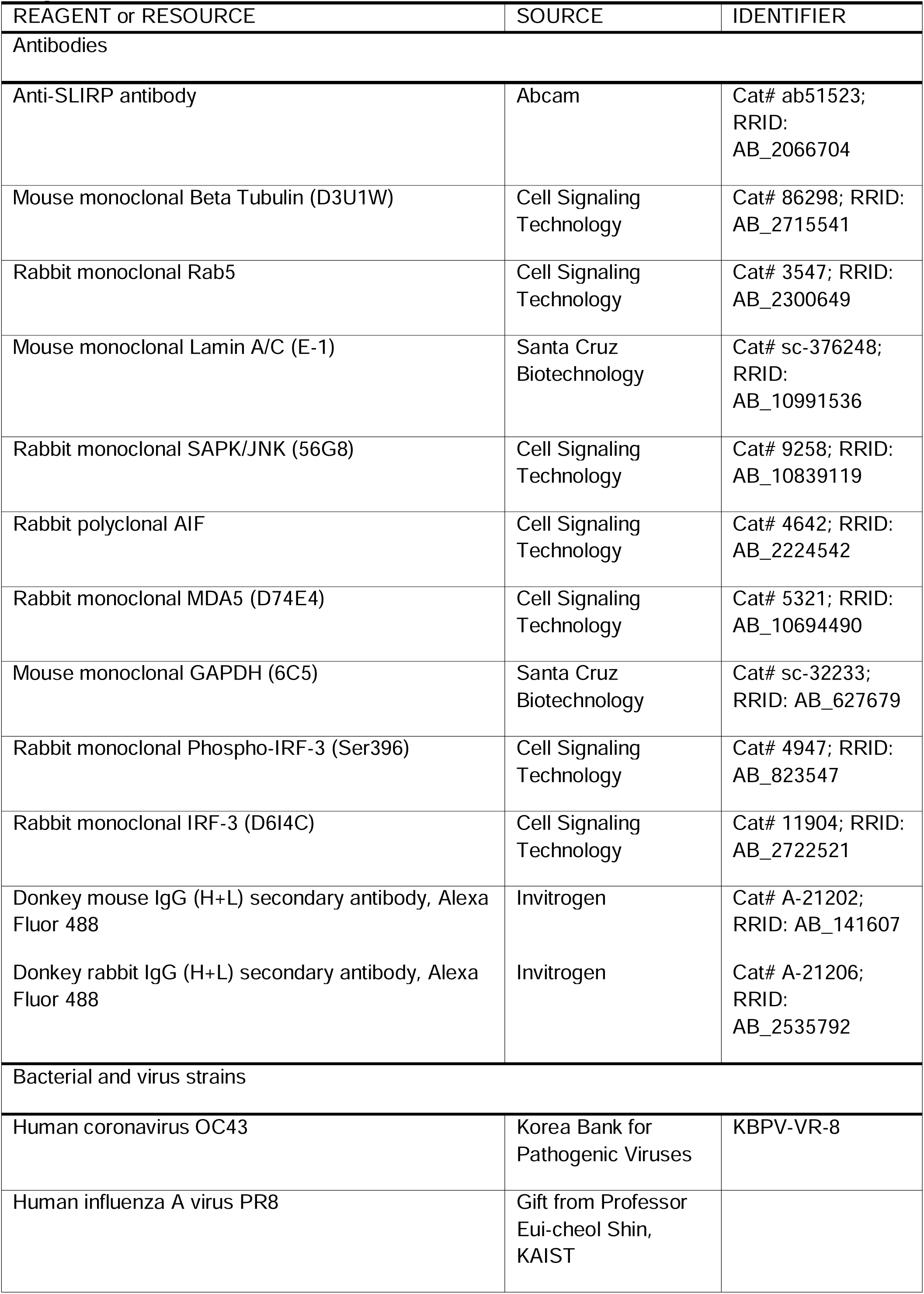
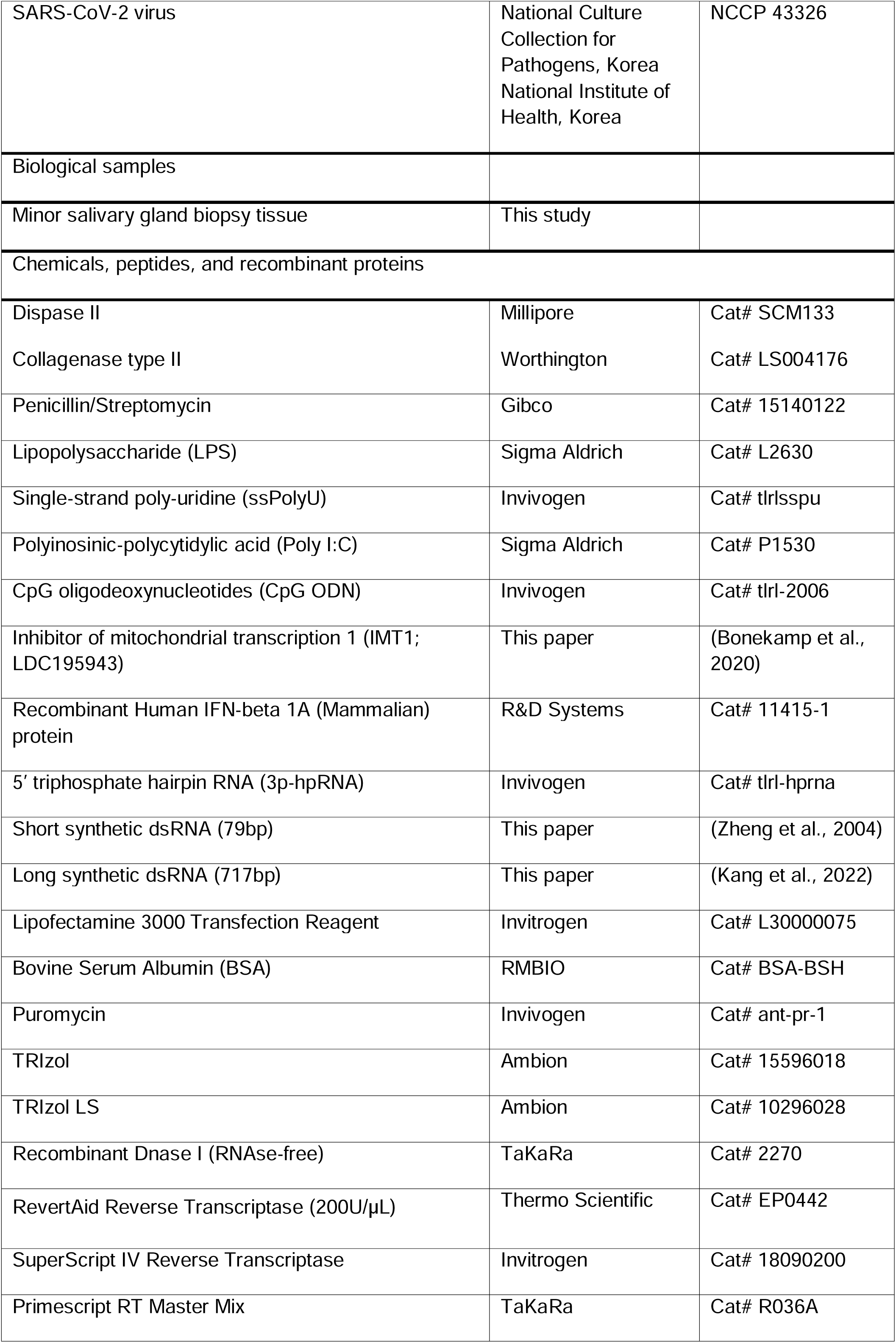
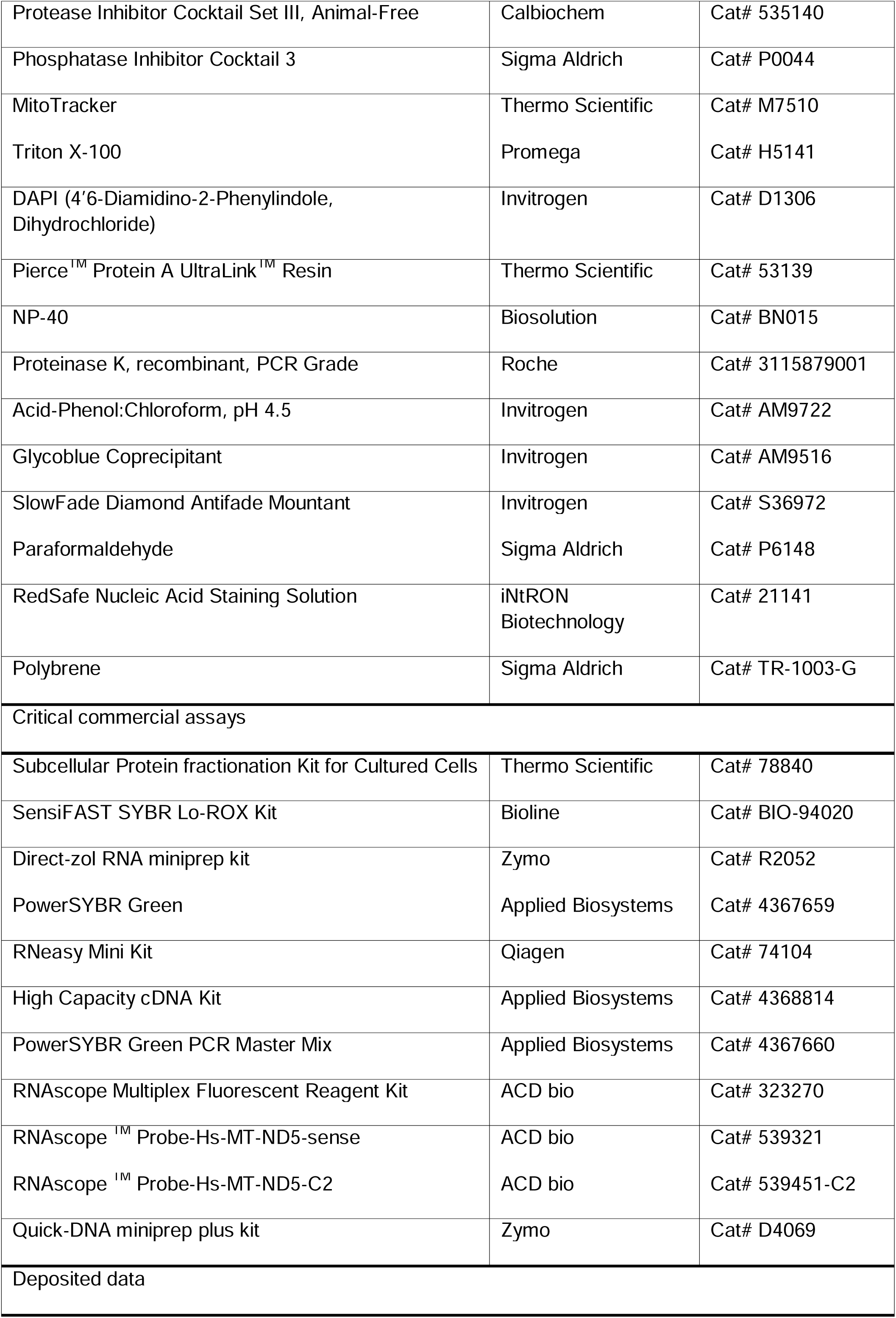
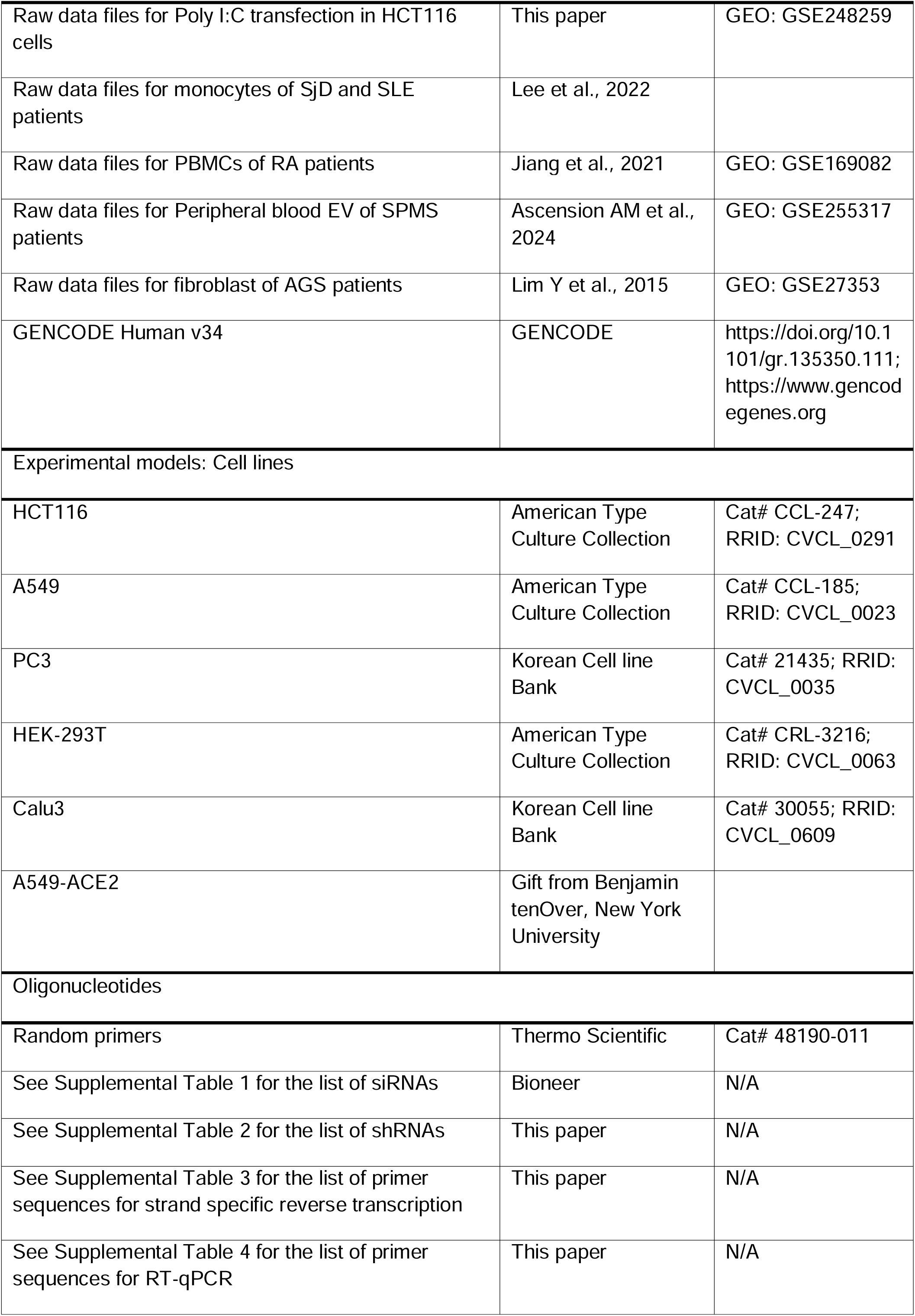
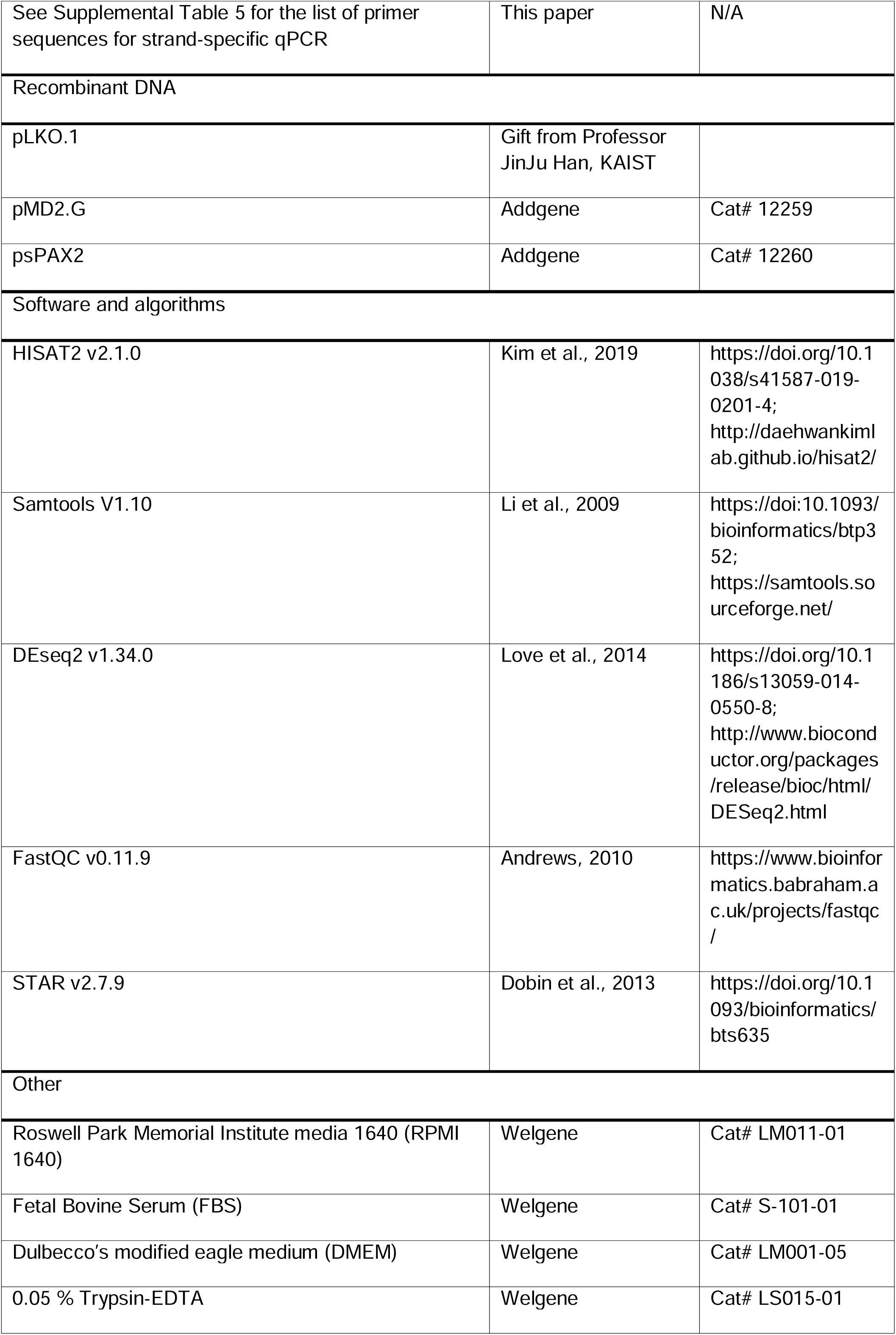
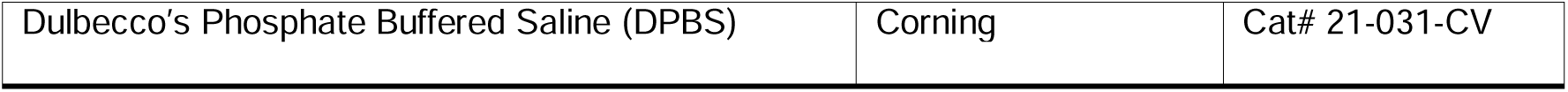

## Resource availability

### Lead contact

Further information and requests for resources and reagents should be directed to and will be fulfilled by the lead contact, Yoosik Kim.

### Materials availability

All reagents and resources used for this study are available upon request to the lead contact.

### Data and code availability

All RNA-seq data have been deposited in GEO and are publicly available as of the date of publication. Accession numbers are listed in the key resources table. The DOI is listed in the key resources table. This paper does not report the original code. No additional information required to reanalyze the data reported in this paper is available from the lead contact upon request.

## Experimental model and study participant details

### Cell lines and maintenance

HCT116 (human colorectal carcinoma cell line), A549 (human lung adenocarcinoma cell line), and PC3 (human prostatic adenocarcinoma cell line) cells were grown in Roswell Park Memorial Institute media 1640 (RPMI 1640) supplemented with 10% (v/v) heat-inactivated fetal bovine serum (FBS). HEK-293T (human embryonic kidney cell line), Calu3 (human lung adenocarcinoma cell line), and A549-ACE2 cells were grown in Dulbecco’s modified eagle medium (DMEM) supplemented with 10% (v/v) FBS. All cells were purchased from the Korean Cell Line Bank while A549-ACE2 was a gift from Benjamin tenOver at New York University. All cells were maintained at 37 °C in a humidified 5% CO2 atmosphere.

For human salivary gland primary cells, the published protocol ^87^ was used with a minor modification. The minor salivary gland biopsy tissue samples were obtained and minced into small pieces in a petri dish with ice-cold Dulbecco’s Phosphate Buffered Saline (DPBS), followed by centrifugation at 400 × g for 5 min and removal of the supernatant. The minced tissues were then resuspended in dispase II, containing 1.5 mg/ml of collagenase type II, and incubated for 30 min at 37 °C in a rotating incubator. After enzymatic digestion, the tissues were disrupted by pipetting, followed by incubation for another 30 min. The digested tissues were passed through a 70 µm cell strainer and washed once with DPBS. The cells were then resuspended in DMEM, containing 10% FBS and 1% Penicillin/Streptomycin, placed in a culture flask, and incubated at 37 °C in a humidified 5% CO2 incubator. Primary cells formed a confluent monolayer within 14 days with fresh media replaced every 4 days. Experiments were performed with the cells between the passages of two and four. The biopsy tissues were obtained from patients at the Center for Orphaned Autoimmune Disorders at the University of Florida College of Dentistry. The informed consent form was signed by each participant according to the protocol (#IRB201900645) approved by the UF Institutional Review Board.

## Method details

### Chemical treatment

Different agonists were directly treated onto cell culture media to stimulate TLRs. Lipopolysaccharide (LPS) was treated with the final concentration of 10 μg/mL, single-strand poly-uridine (ssPolyU) was treated at 5 μg/mL concentration, polyinosinic-polycyticylic acid (poly I:C) was treated at 30 μg/mL, and CpG oligodeoxynucleotides (CpG ODN) was treated at 2 μM. All agonists were treated for 7 h before cell harvest. For the control groups, the same volume of Diethylpyrocarbonate (DEPC)-treated water or DMSO was used. To inhibit mtRNA transcription, a small-molecule inhibitor of mitochondrial RNA polymerase (POLRMT), inositol 4-methyltransferase (IMT1) was treated with the final concentration of 1 μM. To stimulate antiviral signaling, 1000 U of IFN-β was treated for 24 h.

### Transfection

Various agonists were transfected into cells to stimulate cytosolic PRRs. 5′ triphosphate hairpin RNA (3p-hpRNA) was transfected with the final concentration of 0.1 μg/mL, while 1 μg/mL of short synthetic dsRNA, 2 μg/mL of long synthetic dsRNA, 10 μg/mL of poly I:C, or 1.5 μg/mL of CpG ODN was transfected for 7 h prior to cell harvest. Short synthetic dsRNAs with random sequences and a length of 79 base pairs (bp) were prepared as described in a previous study from the Bevilacqua group^88^. Long synthetic dsRNAs were synthesized by in vitro transcribing the *EGFP* gene (717 bp) in sense and antisense direction and annealing the two complementary RNAs, as explained in a previous study from our group^89^. To downregulate gene expressions, predesigned siRNAs were purchased from Bioneer and transfected with the final concentration of 60 nM. After 41 h from siRNA transfection, 1 μg/mL of poly I:C was transfected into cells for 7 h to stimulate cytosolic PRRs without excessive stress. Sequences of the siRNAs used in this study are provided in Table S1. Lipofectamine 3000 Transfection Reagent was used for all transfection.

### Virus infection

A549 cells were washed three times with phosphate-buffered saline (PBS) and infected with human coronavirus OC43 or human influenza A virus PR8 with MOI 1 in the serum-free media. Media was distributed well every 10 min to increase the infection rate for 1 h. Infected cells were washed three times again with PBS and grown in culture media for 24 h at 37 °C in a humidified 5% CO2 atmosphere. Calu3 and A549-ACE2 cells were infected with the SARS-Cov-2 virus with MOI 0.01, 0.1, and 1 for 24 h at 37°C in a humidified 5% CO2 incubator.

### Plasmid subcloning and transduction

To construct shSLIRP-lentiviral particle, shSLIRP and shscramble oligos were designed as described in Supplemental Table 2 and subcloned into pLKO.1 (kindly provided by Professor Jinju Han). Lentiviral particles were produced in HEK-293T cells transfected with pLKO.1-shSLIRP or pLKO.1-shscramble, psPAX2 (Addgene, #12260), and pMD2.G (Addgene, #12259), and supplied by 1% (v/v) Bovine Serum Albumin (BSA). Media containing lentiviral particles were harvested 60 h post-infection and filtered with a 0.45 μm pore filter. Cells were then transduced with lentiviral particles containing shSLIRP or shscramble for 24 h, followed by selection using 5 μg/mL of puromycin containing media for 96 h. Experiments were performed after 48 h of recovery from puromycin selection.

### RNA isolation and RT-qPCR

Total RNAs were isolated directly from cells using 1 ml of TRIzol. For cytosolic RNAs, the Subcellular Protein Fractionation Kit for Cultured Cells was used following the manufacturer’s guide. TRIzol LS reagent was added at a 3:1 ratio to the isolated cytosolic and membrane fraction. Chloroform was added to the TRIzol and nucleic acid was separated. DNase I was treated to remove DNAs and purified RNAs were reverse transcribed using RevertAid reverse transcriptase. For strand-specific RT-qPCR, reverse transcription primers containing CMV promoter were designed to target the specific mtRNAs and SuperScript IV Reverse Transcriptase was used to synthesize the cDNA. qPCR was performed using SensiFAST SYBR Lo-ROX Kit and AriaMX Real-time PCR system.

For the RNAs from SARS-Cov-2 infected cells, cells were treated with TRIzol reagent. RNAs were isolated using the Direct-zol RNA miniprep kit and reverse transcribed using Primescript RT Master Mix. qPCR was performed with PowerSYBR Green and analyzed with QuantStudio 5.

For human minor salivary gland primary cells, total RNA was isolated using RNeasy Mini Kit, followed by reverse transcription to synthesize the first-strand cDNA using High Capacity cDNA Kit. The synthesized cDNA was subjected to quantitative RT-PCR in a 20 μl reaction volume containing Power SYBR Green PCR

Master Mix. RT-qPCR was performed using Applied Biosystems StepOne Real-Time PCR System. Primers used in this study are provided in Table S3-S5.

### Western blotting

Cell lysates were prepared by disrupting cells through sonication in the lysis buffer (50mM Tris-HCl pH 8,0, 100 mM KCl, 0.5% NP-40, 10% Glycerol, and 1 mM DTT) supplemented with a Protease inhibitor cocktail set III. Protein samples were separated on a 10% or 12.5% SDS-PAGE gel and transferred to a PVDF membrane using an Amersham semi-dry transfer system. The membrane was blocked in 5% BSA and incubated with primary antibodies diluted in 1% BSA at a 1:1000 dilution overnight at 4°C. The membrane was incubated with proper secondary antibodies diluted in 1% BSA at a dilution of 1:5000 for 1 h at RT, followed by protein detection with Clarity Western ECL Substrate using ChemiDoc MP Imaging system (Bio-rad).

### Single-molecule RNA fluorescent in situ hybridization

To analyze the expression of heavy and light strands of *ND5* transcript, smRNA-FISH was used. Cells were fixed in 4% (w/v) paraformaldehyde (PFA) for 30 min at room temperature (RT) and sequentially dehydrated with increasing concentrations of ethanol (50%, 70%, and 100%) in PBS for 5 min each. Cells were rehydrated by incubating in EtOH, with decreasing concentrations of ethanol (70%, 50%, and 0%) in PBS for 2 min each, followed by hydrogen peroxide and protease III treatment for 10 min at RT. Pre-treated cells were hybridized with RNAscope RNA-FISH probes and the signal was amplified following the manufacturer’s instructions. Zeiss LSM 780 confocal microscope with a 40X objective (NA= 1.20) was used for visualization and analysis.

### Immunocytochemistry

Cells were prepared in a 0.1% (w/v) gelatin-coated confocal dish (SPL) overnight. To label mitochondria, 80 nM of MitoTracker were added to media and incubated at 37°C in a humidified 5% CO2 atmosphere for 20 min. Cells were fixed using 4% (w/v) PFA for 20 min at RT and permeabilized in 0.1% (v/v/) Triton X-100 diluted in PBS for 15 min. Cells were blocked in 1% (w/v) BSA for 1 h at RT and SLIRP primary antibody was added at a dilution of 1:1000, followed by Alexa fluor fluorophore-labeled secondary antibody incubation at a dilution of 1:1000. 450 nM of 4′5-Diamidino-2- Phenylindole Dihydrochloride (DAPI) was added to counterstain the nuclei. Stained cells were imaged with a Zeiss LSM 780 confocal microscope with a 63X objective (NA= 1.40).

### SLIRP fCLIP

Protein A beads were incubated with SLIRP antibody in the fCLIP lysis buffer (20 mM Tris-HCl pH 7.5, 15 mM EDTA, 0.5% NP-40, 0.1% Triton X-100, 0.1% sodium dodecyl-sulfate (SDS), and 0.1% sodium deoxycholate) for 3 h at 4°C, to prepare SLIRP antibody-conjugated beads. Cells were harvested and fixed with 0.1% (w/v) filtered PFA for 10 min at RT and quenched by adding glycine concentration to the final concentration of 250 mM. The crosslinked cells were lysed by incubating for 10 min on ice and then sonicated for complete lysis. The lysate was incubated in the SLIRP antibody-conjugated beads for 3 h at 4°C. The beads are washed 4 times with the fCLIP wash buffer (20 mM Tris-HCl pH 7.5, 150 mM NaCl, 10mM EDTA, 0.1% NP-40, 0.1% SDS, 0.1% Triton X-100, and 0.1% sodium deoxycholate). SLIRP-RNA complex was eluted from the beads by incubating in elution buffer (200 nM Tris-HCl pH7.4, 100 mM NaCl, 20 mM EDTA, 2% SDS, and 7M Urea) for 3 h at 25 °C. The eluate was treated with 2 mg/mL of proteinase K overnight at 65°C. RNA was purified using acid-phenol:Chloroform pH 4.5 extraction.

### DNA purification, qPCR, and agarose gel electrophoresis

Cellular DNA was isolated using Quick-DNA miniprep plus kit following manufacturer’s instructions. qPCR was performed as described previously. Amplified DNAs were separated on a 2% agarose gel and Redsafe Nucleic Acid Staining Solution was used for visualization.

### Statistical and functional analysis of mRNA-seq data

For the analysis of RNA-seq data after poly I:C transfection into HCT116, sequencing read files were quality-checked with FastQC (version 0.11.9) for any abnormalities. Reads were aligned against the reference human genome (GRCh38) with STAR aligner (version 2.7.9) ^90^. Gene quantification was performed simultaneously by providing --quantMode GeneCounts option. Reads were quantified against GENCODE human genome annotation (version 38) ^91^. Differential gene expression analysis and other statistical calculations were performed using the DESeq2 package in R ^92^.

For the analysis of RNA-seq data of SjD, SLE, RA, SPMS, and AGS patients, raw reads were mapped to the reference human genome (GRCh38) using HISAT2 v.2.1.0 using default parameters ^93^. Uniquely and concordantly mapped reads were selected using samtools and then quantified against the GENCODE human genome (version 34) using featureCounts. Differential gene expression analysis and other statistical calculations were performed using the DESeq2 package in R.

### Quantification and statistical analysis

Continuous variables were analyzed using the one-tailed Student’s t-test. All data were biologically replicated at least three times. The error bars indicate the standard error of the mean. P values ≤0.05 were considered statistically significant (∗p ≤ 0.05, ∗∗p ≤ 0.01, and ∗∗∗p ≤ 0.001).

## Notes

### Summary of Updates

The abnormal innate immune response is a prominent feature underlying autoimmune diseases. One emerging factor that can trigger dysregulated immune activation is cytosolic mitochondrial double-stranded RNAs (mt-dsRNAs). However, the mechanism by which mt-dsRNAs stimulate immune responses remains poorly understood. Here, we discover SRA stem-loop interacting RNA binding protein (SLIRP) as a key amplifier of mt-dsRNA-triggered antiviral signals. In autoimmune diseases, SLIRP is commonly upregulated, and targeted knockdown of SLIRP dampens the interferon response. We find that the activation of melanoma differentiation-associated gene 5 (MDA5) by exogenous dsRNAs upregulates SLIRP, which then stabilizes mt-dsRNAs and promotes their cytosolic release to activate MDA5 further, augmenting the interferon response. Furthermore, the downregulation of SLIRP partially rescues the abnormal interferon-stimulated gene expression in autoimmune patients primary cells and makes cells vulnerable to certain viral infections. Our study unveils SLIRP as a pivotal mediator of interferon response through positive feedback amplification of antiviral signaling.

