## Supplemental informations for "SLIRP promotes autoimmune diseases by amplifying antiviral signaling via positive feedback regulation"

**Supplemental Information**

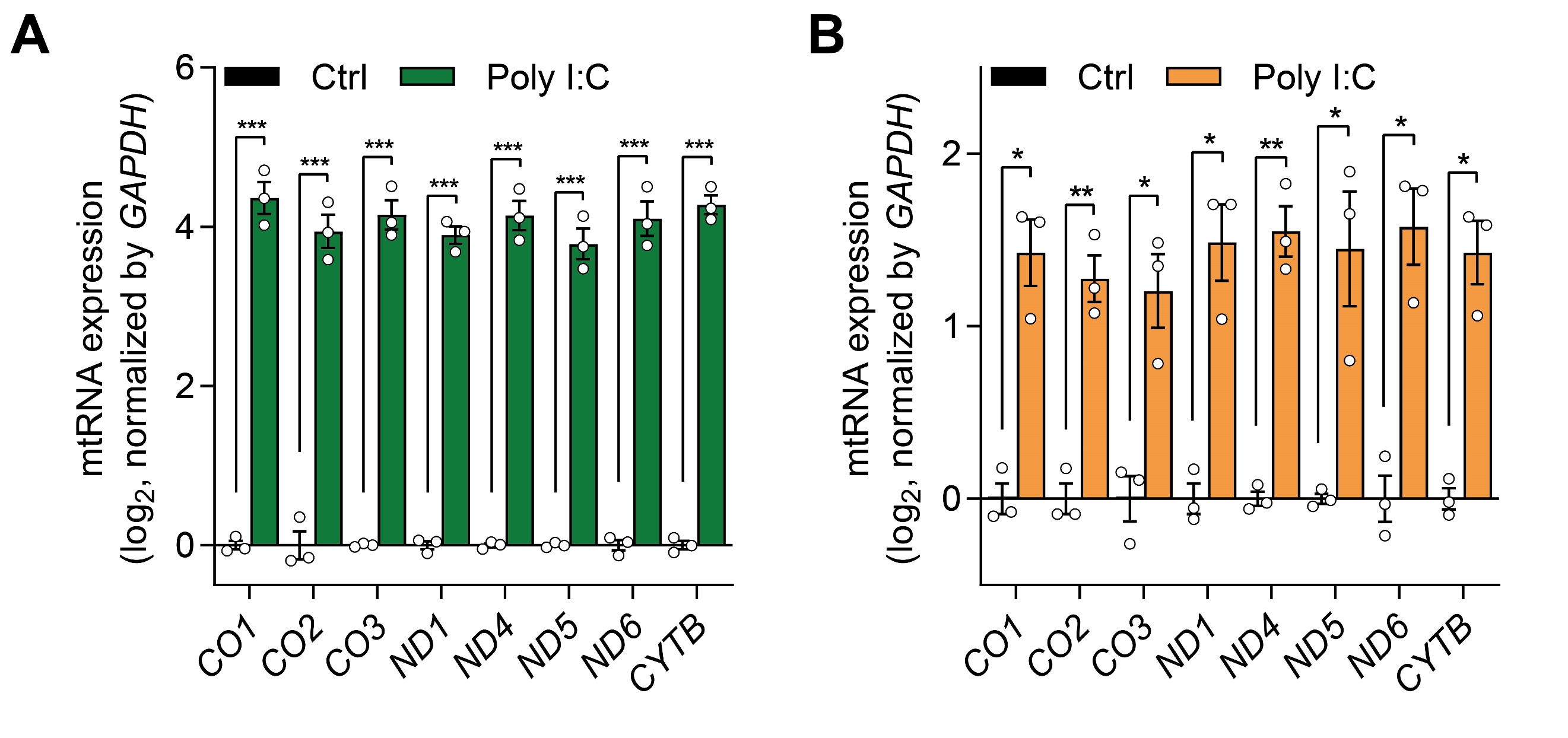

**Figure S1. Effect of Poly I:C transfection on mtRNA expression.** (A, B) Analysis of total mtRNA expression upon poly I:C transfection in A549 (A) and PC3 (B). The average of three biological replicates are shown and error bars denote s.e.m. All of the statistical significances were calculated using one-tail Student’s t-tests; * p ≤ 0.05, ** p ≤ 0.01, and *** p ≤ 0.001.

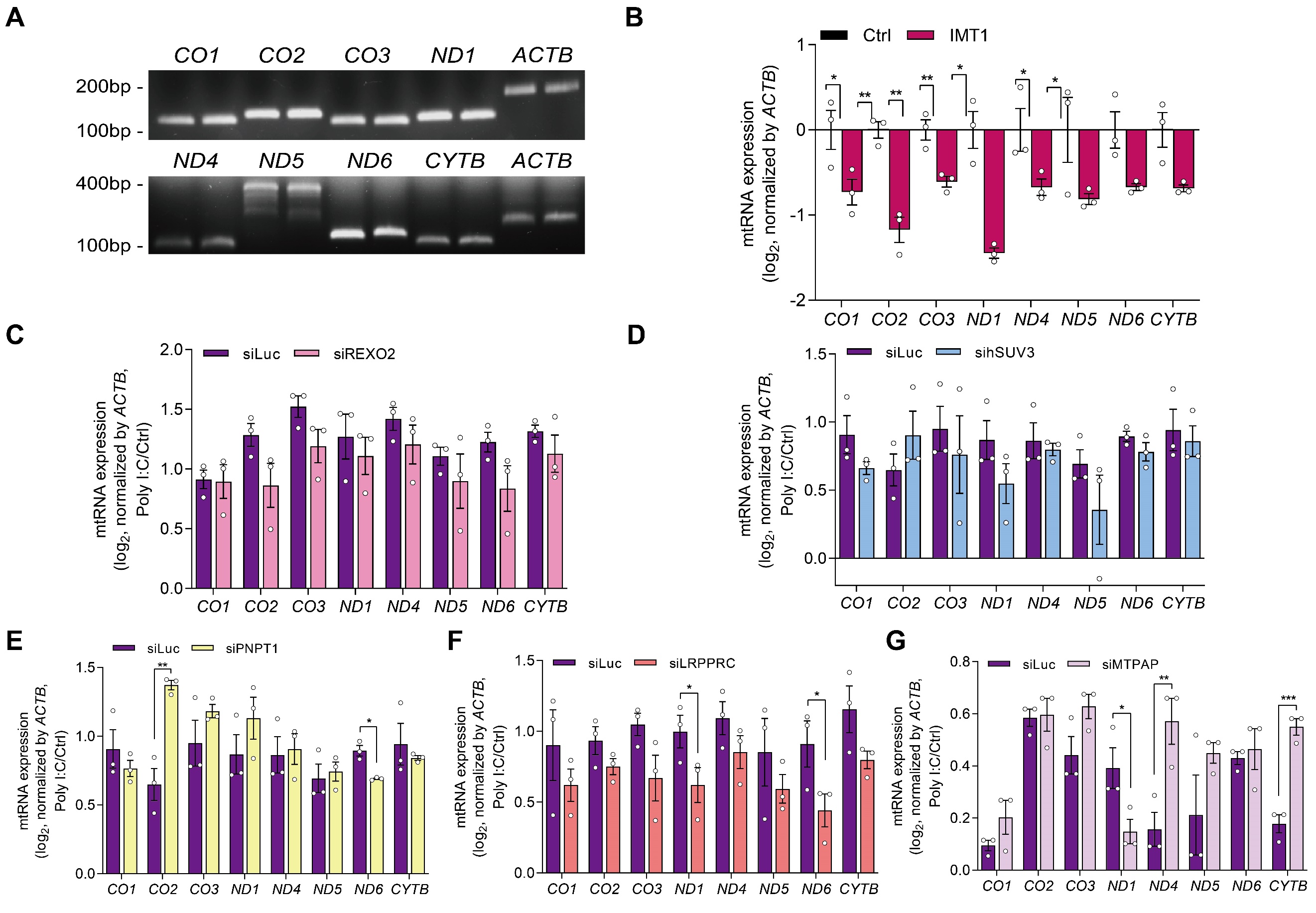

**Figure S2. Downregulation of mtRNA stability regulators on poly I:C mediated mtRNA induction**. (A) Confirmation of the qPCR result by agarose gel electrophoresis to detect mtDNA amplicon. (B) Analysis of total mtRNA expression 48h after IMT1 treatment. (C-G) Analysis of total mtRNA expression before and after poly I :C transfection in control and REXO2 (C), hSUV3 (D), PNPT1 (E), LRPPRC (F), or MTPAP knocked down HCT116 cells (G). Data are normalized only to those of DEPC-transfected cells in order to analyze the degree of mtRNA induction by poly I :C. Three independent experiments were carried out, and error bars denote s.e.m. All of the statistical significances were calculated using one-tail Student’s t-tests; * p ≤ 0.05, ** p ≤ 0.01, and *** p ≤ 0.001.

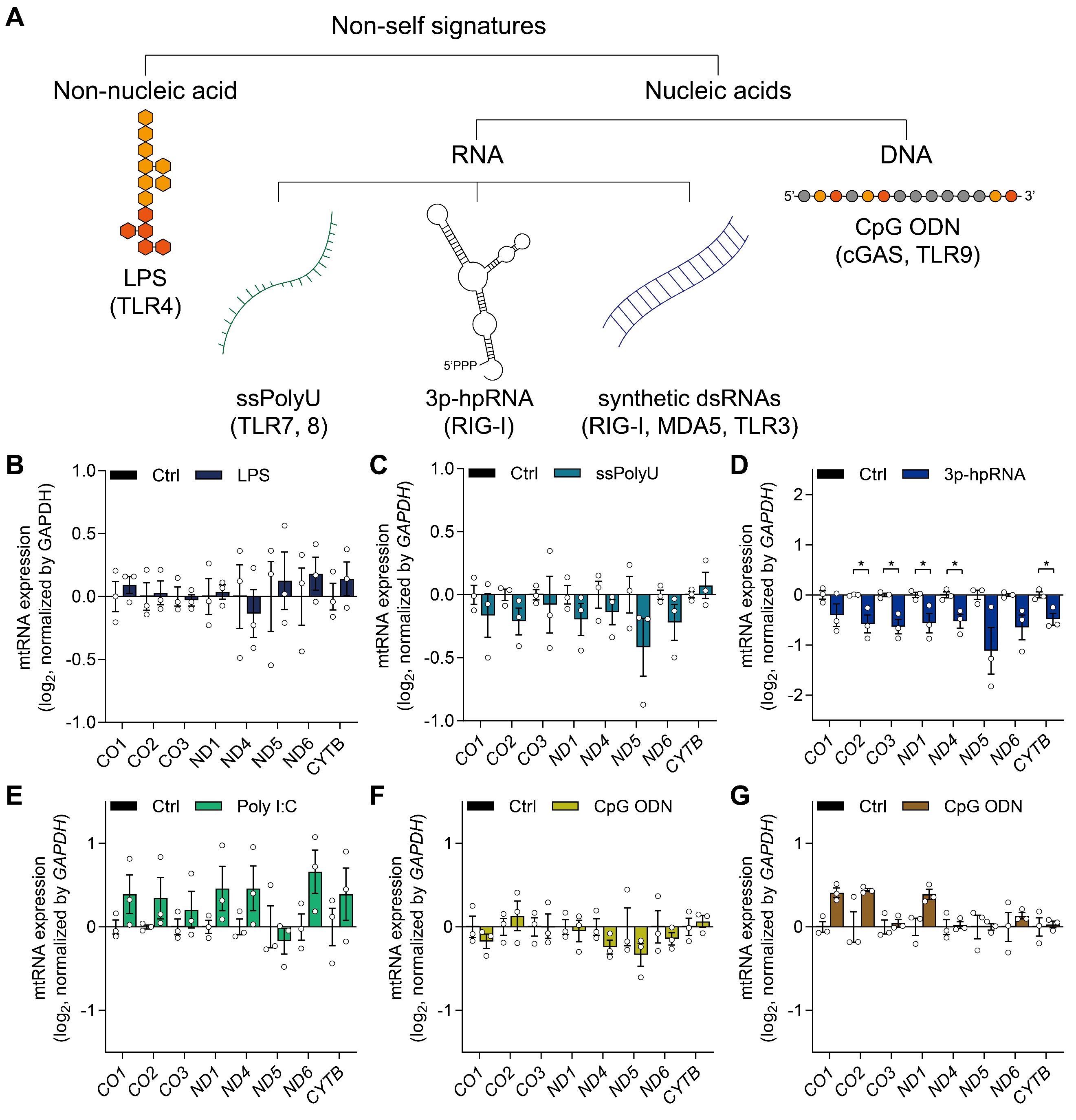

**Figure S3. Effect of PRR agonists on mtRNA expression.** (A) A scheme for the immune stressors represents various non-self signatures and the corresponding PRRs. (B-G) Analysis of total mtRNA expression upon LPS treatment (B), ssPolyU treatment (C), 3p-hpRNA transfection (D), poly I:C treatment (E), CpG ODN treatment (F), and CpG transfection (G). N=3 and error bars denote s.e.m. All of the statistical significances were calculated using one-tail Student’s t-tests; * p ≤ 0.05, ** p ≤ 0.01, and *** p ≤ 0.001.

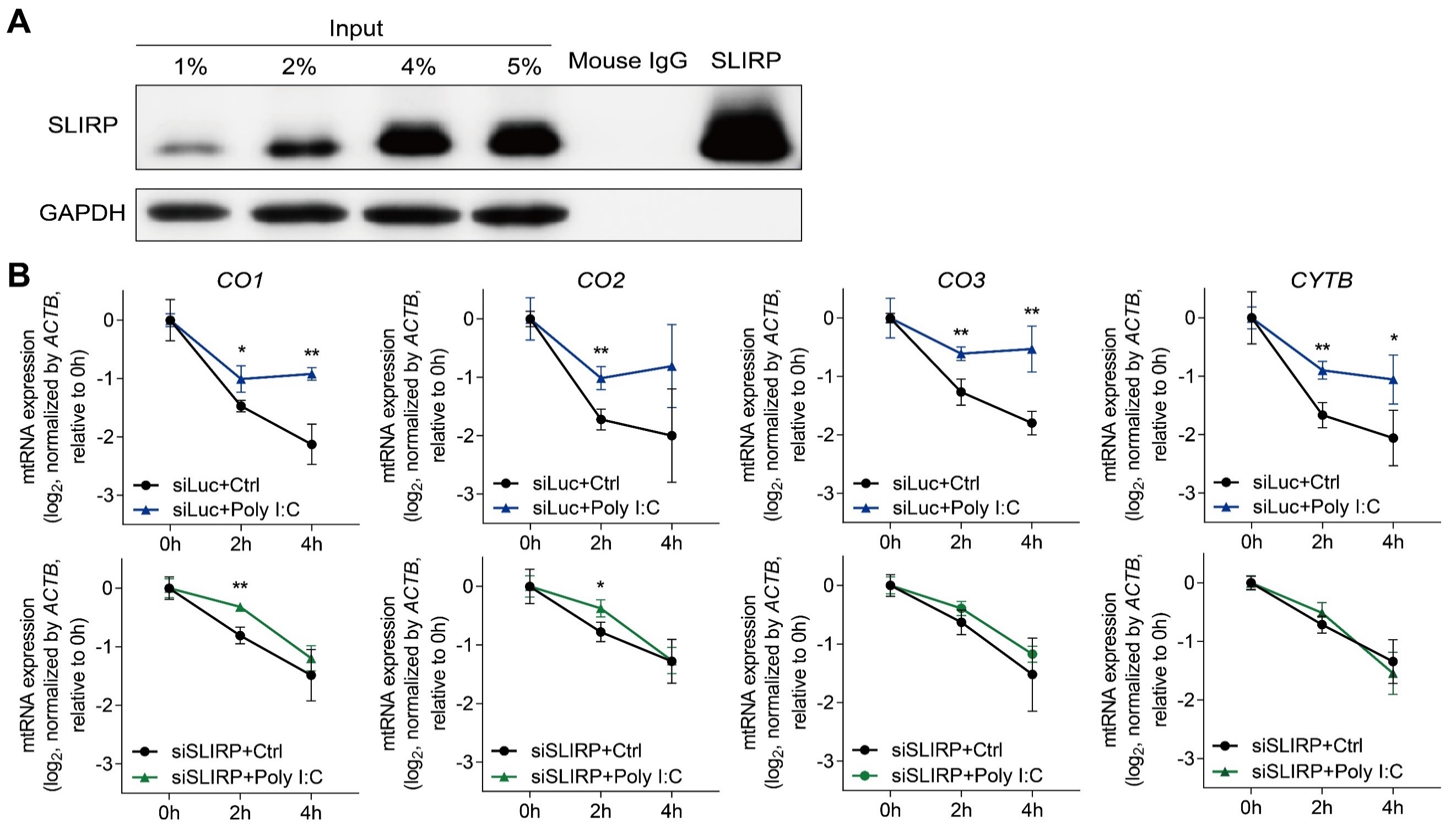

**Figure S4. Stabilization of mtRNAs by SLIRP.** (A) Analysis of protein expression of input and SLIRP fCLIP. SLIRP antibody specifically interact with SLIRP protein, while mouse IgG did not. (B) Analysis of mtRNA stability before and after poly I :C transfection in control and SLIRP-deficient cells. Three independent experiments were carried out, and error bars denote s.e.m. All of the statistical significances were calculated using one-tail Student’s t-tests; * p ≤ 0.05, ** p ≤ 0.01, and *** p ≤ 0.001.

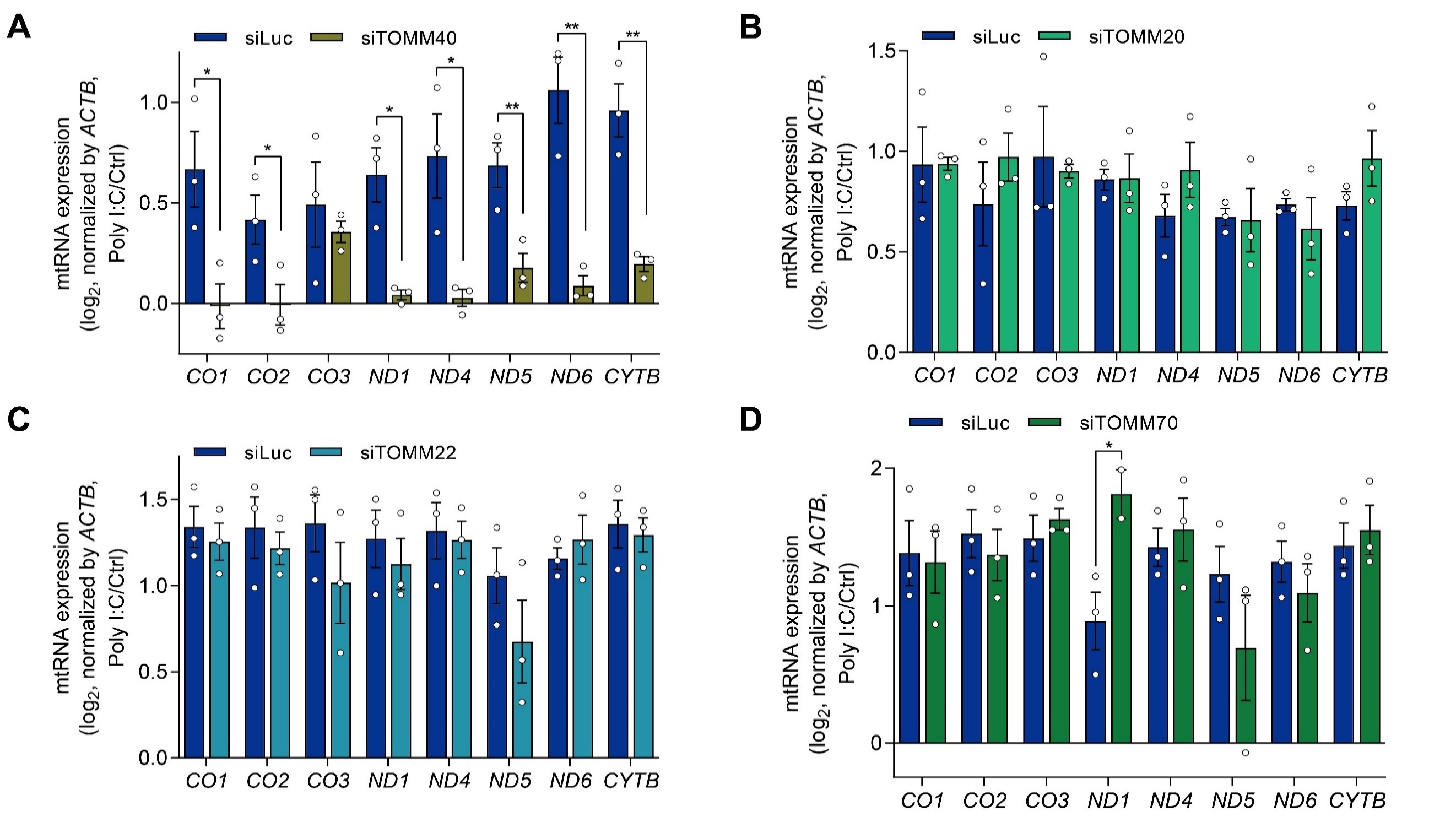

**Figure S5. Downregulation of components of the TOM complex on poly I:C-mediated mtRNA induction.** (A-D) Analysis of total mtRNAs upon poly I :C in control and TOMM40 (A), TOMM20 (B), TOMM22 (C), and TOMM70 knocked down HCT116 cells (D). Three independent experiments were carried out, and error bars denote s.e.m. All of the statistical significances were calculated using one-tail Student’s t-tests; * p ≤ 0.05, ** p ≤ 0.01, and *** p ≤ 0.001.

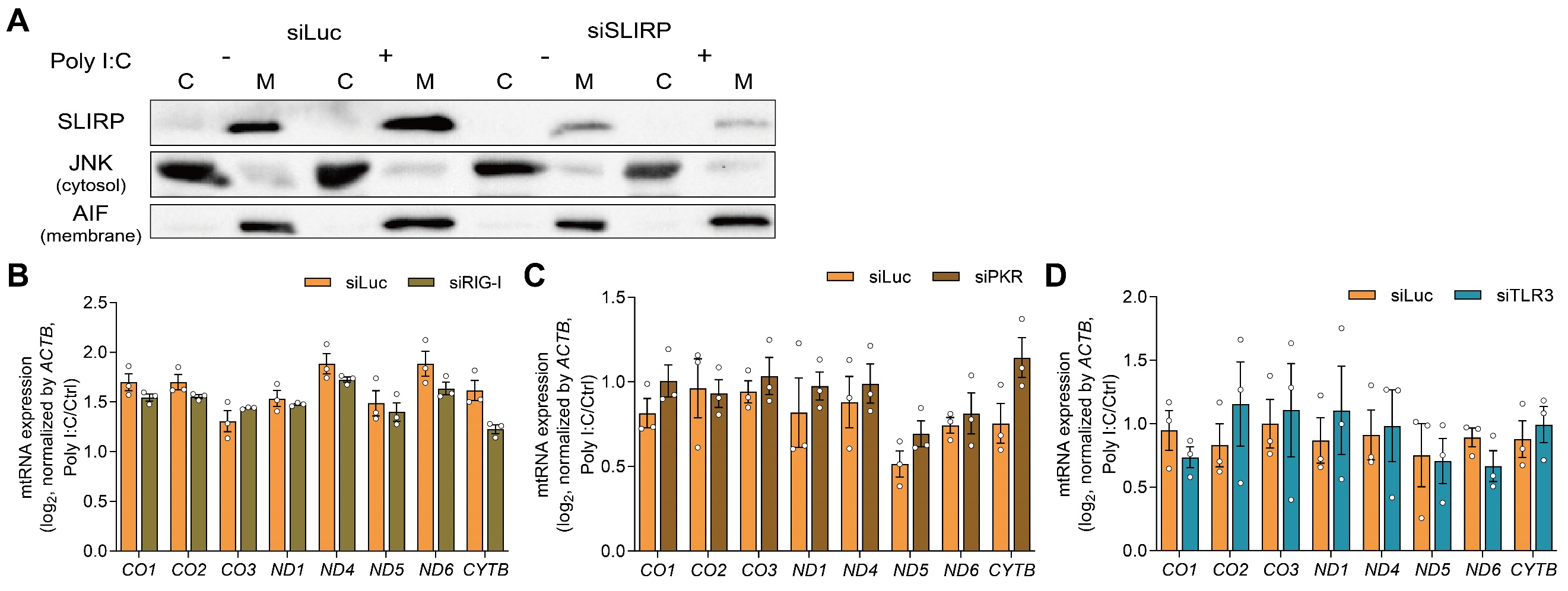

**Figure S6. Downregulation of dsRNA sensors on poly I:C-mediated mtRNA induction.** (A) Western blot analysis for SLIRP expression upon poly I :C in the cytosol and membrane fraction, with or without SLIRP. JNK is used as a marker for the cytosol fraction while AIF is used as a marker of the membrane fraction. (B-D) The effect of RIG-I (B), PKR (C), and TLR3 (D) downregulation on mtRNA induction by poly I :C. Three independent experiments were carried out, and error bars denote s.e.m. All of the statistical significances were calculated using one-tail Student’s t-tests; * p ≤ 0.05, ** p ≤ 0.01, and *** p ≤ 0.001.

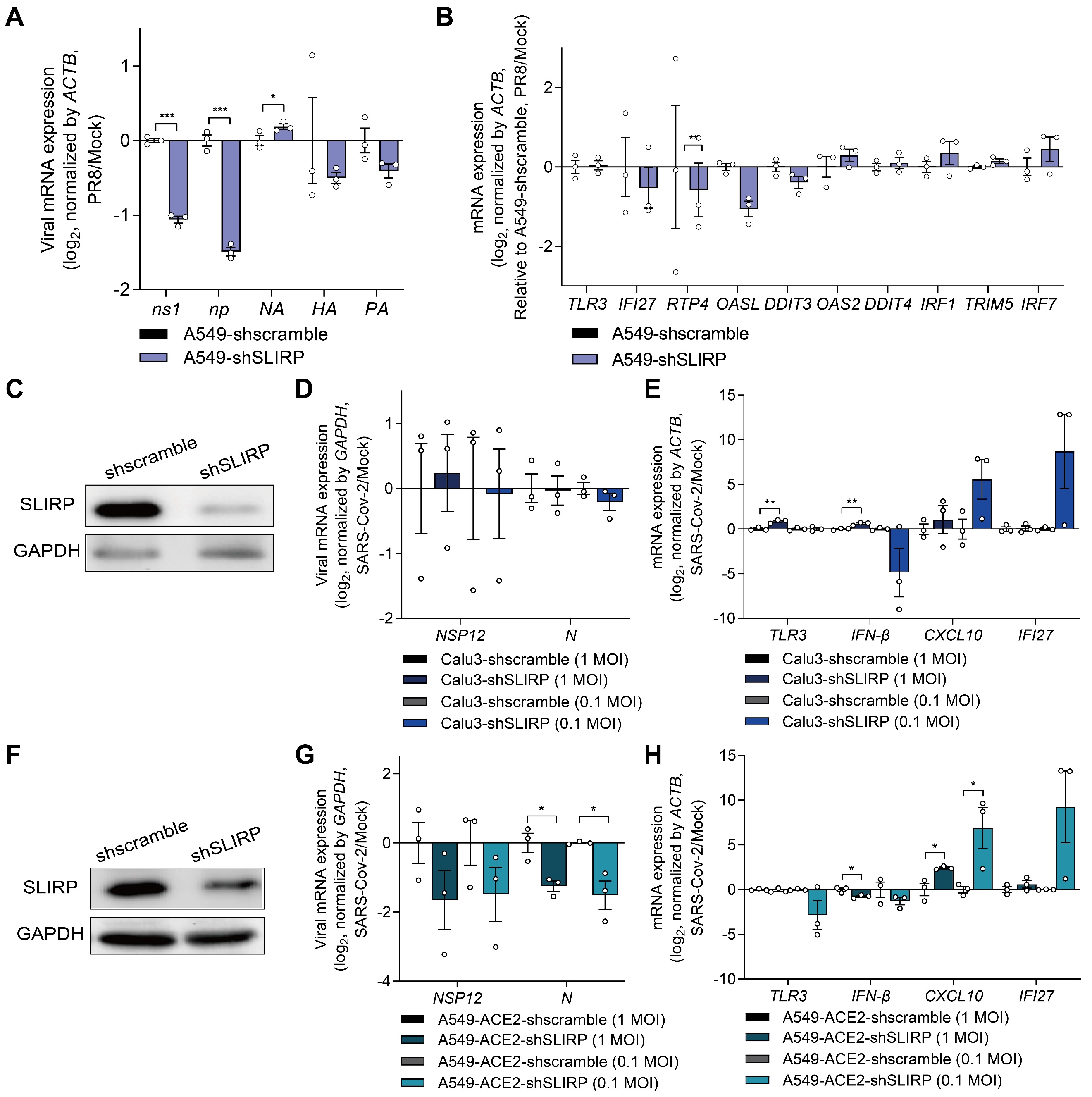

**Figure S7. Downregulation of SLIRP on viral replication and ISG induction for PR8 and SARS-CoV-2.** (A, B) Analysis of viral mRNA expression (B) and ISG induction (C) after infecting SLIRP-deficient A549 with PR8 at MOI=1. (C) Western blot analysis of SLIRP expression in Calu3 cells transduced with shSLIRP. (D, E) Analysis of viral mRNA expression (D) and ISG induction (E) after infecting SLIRP-deficient Calu3 with SARS-CoV-2 at MOI=0.1, 1. (F) Western blot analysis of SLIRP expression in A549-ACE2 cells transduced with shSLIRP. (D, E) Analysis of viral mRNA expression (D) and ISG induction (E) after infecting SLIRP-deficient A549-ACE2 with SARS-CoV-2 at MOI=0.1 and 1. Three independent experiments were carried out, and error bars denote s.e.m. All of the statistical significances were calculated using one-tail Student’s t-tests; * p ≤ 0.05, ** p ≤ 0.01, and *** p ≤ 0.001.

**Table S1**. Sequences of siRNAs

| **Gene** | **Sequences (5’-3’)** |
| --- | --- |
| siLuc | CUU ACG CUG AGU ACU UCG A |
| siSLIRP | CAG UUC GGC CAU GUC AGA A |
| siLRPPRC | CUG AAC GAU GCU GCC AAC A |
| siMTPAP | CAG AUC UGC ACA CAC UGA A |
| siREXO2 | CAG GAU UGG ACA UUG AGA A |
| sihSUV3 | CUG CUA UUG ACC UGG UGA U |
| siPNPT1 | GUA AGU UGU GAG GUA GAU A |
| siTOMM40 | CUG UAC UGG UGG GUG ACA U |
| siTOMM20 | GUG UCA UCC UAA GUA CCU U |
| siTOMM22 | GUG ACA UAU UCU CCG UAG U |
| siTOMM70 | CAC UCU AAU GGA UAG UGU A |
| siBAK1 | CUC AAG AGU ACA GAA GCU U |
| siMDA5 | GAA AAU GCA UCA CGU CAA U |
| siRIG-I | CUC UUG AUG CGU CAG UGA U |
| siPKR-1 | GCA GGG AGU AGU ACU UAA A |
| siPKR-2 | GCA UGG GCC AGA AGG AUU U |
| siPKR-3 | GCA GAU ACA UCA GAG AUA A |
| siTLR3 | ACU UUG CCU UGU AUC UAC U |

**Table S2**. Primer sequences for shRNAs

| **Gene** | **Forward Primer (5’-3’)** | **Reverse Primer (5’-3’)** |
| --- | --- | --- |
| shscramble | CCGGTCCTAAGGTTAAGTCGCCCTCGCTCGAGCGAGGGCGACTTAACCTTAGGTTTTTG | TCCTAAGGTTAAGTCGCCCTCGCTCGAGCGAGGGCGACTTAACCTTAGGTTTTTGAATT |
| shSLIRP | CCGGGCGCTGCGTAGAAGTATCAATCTCGAGATTGATACTTCTACGCAGCGCTTTTTG | AATTCAAAAAGCGCTGCGTAGAAGTATCAATCTCGAGATTGATACTTCTACGCAGCGC |

**Table S3**. Primer sequences for strand-specific reverse transcription

| **Gene** | **Sequences (5’-3’)** |
| --- | --- |
| ND1 Heavy | CGCAAATGGGCGGTAGGCGTGGTTGTGATAAGGGTGGAGAGG |
| ND1 Light | CGCAAATGGGCGGTAGGCGTGTCAAACTCAAACTACGCCCTG |
| ND4 Heavy | CGCAAATGGGCGGTAGGCGTGTGTTTGTCGTAGGCAGATGG |
| ND4 Light | CGCAAATGGGCGGTAGGCGTGCCTCACACTCATTCTCAACCC |
| ND5 Heavy | CGCAAATGGGCGGTAGGCGTGTTTGGGTTGAGGTGATGATG |
| ND5 Light | CGCAAATGGGCGGTAGGCGTGCATTGTCGCATCCACCTTTA |
| ND6 Heavy | CGCAAATGGGCGGTAGGCGTGGGTTGAGGTCTTGGTGAGTG |
| ND6 Light | CGCAAATGGGCGGTAGGCGTGCCCATAATCATACAAAGCCCC |
| CYTB Heavy | CGCAAATGGGCGGTAGGCGTGGGATAGTAATAGGGCAAGGACG |
| CYTB Light | CGCAAATGGGCGGTAGGCGTGCAATTATACCCTAGCCAACCCC |
| GAPDH | CGCAAATGGGCGGTAGGCGTGTGAGCGATGTGGCTCGGCT |
| ACTB | CGCAAATGGGCGGTAGGCGTGACA CAG AGTACTTGCGCTCAG |

**Table S4**. Primer sequences for RT-qPCR

| **Gene** | **Forward Primer (5’-3’)** | **Reverse Primer (5’-3’)** |
| --- | --- | --- |
| CO1 | GCCATAACCCAATACCAAACG | TTGAGGTTGCGGTCTGTTAG |
| CO2 | CTAGTCCTCTATGCCCTTTTCC | GTAAAGGATGCGTAGGGATGG |
| CO3 | CCTTTTACCACTCCAGCCTAG | CTCCTGATGCGAGTAATACGG |
| ND1 | TCAAACTCAAACTACGCCCTG | GTTGTGATAAGGGTGGAGAGG |
| ND4 | CTCACACTCATTCTCAACCCC | TGTTTGTCGTAGGCAGATGG |
| ND5 | CTAGGCCTTCTTACGAGCC | CGCAAATGGGCGGTAGGCGTGT  TTGGGTTGAGGTGATGATG |
| ND6 | TGCTGTGGGTGAAAGAGTATG | CGCAAATGGGCGGTAGGCGTGC  CCATAATCATACAAAGCCCC |
| CYTB | CAATTATACCCTAGCCAACCCC | GGATAGTAATAGGGCAAGGACG |
| SLIRP | AACACTTTGCACAGTTCGGC | ACCCAACCCAAACCTCTGTG |
| TOMM20 | GTATGCGGGGCCCTTTTCAT | TGAAGTTGGGGTCACTTCGT |
| TOMM22 | TACAGGTTTTCCAGGGCAGC | CCGTCTCAAAGACAACGGGA |
| TOMM40 | TAGCAACTGGATCGTGGGTG | GTGATTCAGGAAGGCCCCAA |
| TOMM5 | AAGATGCGCGAGGATGTGAT | AATGGAGTGACTCGCAGGAG |
| TOMM7 | GCTGGGGCTTTATCCCTCTT | TTGGTTCAGGCATTCCGGG |
| TOMM70 | TGCTGCCTTTGAACAGTTGC | CTCATGGGCTTTTGCACGTC |
| IFIT2 | AGAGCGAAGGTGTGCTTTGA | AGGGTCAATGGCGTTCTGAG |
| IFNB1 | TCTCCTGTTGTGCTTCTCCAC | GCCTCCCATTCAATTGCCAC |
| IFIT1 | AGAGAAAAAGCAGGACCCACAA | CACCATTTGTACACATCTCCACTG |
| IFIT3 | GAAGGAACTGGGCCGCCTGCTAAG | GCCCTGGCCCATTTCCTCACTACC |
| OAS2 | GGTTCACCATCCAGGTGTTCA | AGCAATGCTTACTCAGAGCGT |
| OASL | TTCAGCGAGCTGCAGAGAAA | CCCTCTGCTCCACTGTCAAG |
| MX2 | ACCGAGCTAGAGCTTCAGGA | TCAGGGGAGGTGATCTCCAG |
| MX1 | TTCTGGGTCGGAGGCTACAG | TGGATGGCGGCGTTCT |
| XAF1 | AGCGCAGGAAAGTCAAGACC | TGAGTCTGGACAACATTTACCC |
| ISG20 | GGTGCTGTGCTGTACGACAA | GAGCTGCAGGATCTCTAGCC |
| OAS1 | GAGACCCAAAGGGTTGGAGG | TGTGCTGGGTCAGCAGAATC |
| OAS3 | CAAGGTGGTCAAGGAGCGG | CAGCATCGTCTGGGATGTCA |
| IRF1 | AAGCATGGCTGGGACATCAA | TGCTTTGTATCGGCCTGTGT |
| IRF7 | CTGTGGACACCTGTGACACC | TGCCCTCTCAGGAGCCAA |
| STAT1 | AACCTCGACAGTCTTGGCAC | CACTGAGACATCCTGCCACC |
| ISG15 | GATCACCCAGAAGATCGGCG | GTTCGTCGCATTTGTCCACC |
| IFI6 | GGGTGGAGGCAGGTAAGAAA | GTCAGGGCCTTCCAGAACC |
| IRF2 | GGCTAGACATGGGTGGGATG | GCGCATCTGAAATTCGCCTT |
| SP100 | AGGTGTGCAACAAATGGGGA | ATGCAACTCCACGGGTTCTT |
| USP18 | GGCTCCTGAGGCAAATCTGT | CAACCAGGCCATGAGGGTAG |
| IFI27 | ATCAGCAGTGACCAGTGTGG | TGGCCACAACTCCTCCAATC |
| MDA5 | TTGGACTCGGGAATTCGTGG | AACGATGGAGAGGGCAAGTC |
| NFKBIA | CTCCGAGACTTTCGAGGAAATAC | GCCATTGTAGTTGGTAGCCTTCA |
| IRF3 | ACACATACTGGGCAGTGAGC | CTACAATGAAGGGCCCCAG |
| IFITM1 | CGGCTCTGTGACAGTCTACC | TGCACAGTGGAGTGCAAAGG |
| AMY1A | CCTTCTGGGATGCTAGGCTG | ATCTTGGCCAACGGTAGCTT |
| SLC12A2 | TGGCACCAAGGATGTGGTAG | GGTTGAGTTGCAGTCTTGCC |
| MUC7 | CCACACCTAATTCTTCCC | CTATTGCTCCACCATGTC |
| AQP5 | CCGCTCACTGGGTTTTCTGG | TTTGATGATGGCCACACGCT |
| ns2a  (OC43) | CCCATTTTCAGGGTTTTGTG | CACCCCAATGCATATCATCA |
| ns12.9  (OC43) | CTTGGTGCCGTAATCAAGGT | TGTCTGCGCCAGGTAATAAA |
| ns3  (OC43) | GTGGCATTTTTGGCAACTTT | TCATCCACATCAAGGACTGG |
| N  (OC43) | GCTGCCACGATGGTATTTTT | CGGAATAGCCTCATCGCTAC |
| S_RBD  (OC43) | GCTATACCCAATGGCAGGAA | CTGCAGGTCGAGGCTTAAAA |
| S_2  (OC43) | CGCTAACTCTTCCGAACCAG | ACCAGTGGTAATCGCTCCAC |
| TLR3 | TAGCAGTCATCCAACAGAATCAT | AATCTTCTGAGTTGATTATGGGTAA |
| CXCL10 | AGCAGAGGAACCTCCAGTCT | ATGCAGGTACAGCGTACAGT |
| RTP4 | GACACAGCCAATTGTGAGGC | AGGGATTTGGACGGCTTTGT |
| DDIT3 | GTTCCAGCCACTCCCCATTA | GTCCCGAAGGAGAAAGGCAA |
| DDIT4 | ACACTTGTGTGCCAACCTGA | CAGGCGCAGTAGTTCTTTGC |
| TRIM5 | GACGGAGAACGTGACCTTGAA | CTGTCACATCAACCCAGTAGC |
| ns1 (PR8) | CAGCACTCTTGGTCTGGACA | TGGACCATTCCCTTGACATT |
| np (PR8) | TGCTTCAAAACAGCCAAGTG | GCCCAGTACCTGCTTCTCAG |
| NA (PR8) | GTTGATGGAGCAAACGGAGT | ACCAATCAGTCATTGCCACA |
| HA (PR8) | TTGCTAAAACCCGGAGACAC | CCTGACGTATTTTGGGCACT |
| PA (PR8) | AGCCTATGTGGATGGATTCG | GCATCCATCAGCAGGAATTT |
| NSP12  (SARS-Cov-2) | CAGAGAGCTAGGTGTTGTAC | AAGCACGTAGTGCGTTTATC |
| N  (SARS-Cov-2) | CAGCGTTCTTCGGAATGTCG | AGGCTTGAGTTTCATCAGCC |
| GAPDH | CTCCTCCACCTTTGACGCTG | TCCTCTTGTGCTCTTGCTGG |
| ACTB | CCTGTACGCCAACACAGTGC | ATACTCCTGCTTGCTGATCC |

**Table S5**. Primer sequences for strand-specific qPCR

| **Gene** | **Forward Primer (5’-3’)** | **Reverse Primer (5’-3’)** |
| --- | --- | --- |
| ND1 Heavy | TCAAACTCAAACTACGCCCTG | CGCAAATGGGCGGTAGGCGTG |
| ND1 Light | GTTGTGATAAGGGTGGAGAGG | CGCAAATGGGCGGTAGGCGTG |
| ND4 Heavy | CTCACACTCATTCTCAACCCC | CGCAAATGGGCGGTAGGCGTG |
| ND4 Light | TGTTTGTCGTAGGCAGATGG | CGCAAATGGGCGGTAGGCGTG |
| ND5 Heavy | CTAGGCCTTCTTACGAGCC | CGCAAATGGGCGGTAGGCGTG |
| ND5 Light | TAGGGAGAGCTGGGTTGTTT | CGCAAATGGGCGGTAGGCGTG |
| ND6 Heavy | TCATACTCTTTCACCCACAGC | CGCAAATGGGCGGTAGGCGTG |
| ND6 Light | TGCTGTGGGTGAAAGAGTATG | CGCAAATGGGCGGTAGGCGTG |
| CYTB Heavy | CAATTATACCCTAGCCAACCCC | CGCAAATGGGCGGTAGGCGTG |
| CYTB Light | GGATAGTAATAGGGCAAGGACG | CGCAAATGGGCGGTAGGCGTG |
| GAPDH | CAACGACCACTTTGTCAAGC | CGCAAATGGGCGGTAGGCGTG |
| ACTB | ACACAGTGCTGTCTCGTGGTA | CGCAAATGGGCGGTAGGCGTG |
